## Supplemental figures for "scSemiProfiler: Advancing Large-scale Single-cell Studies through Semi-profiling with Deep Generative Models and Active Learning"

### Supplementary figures

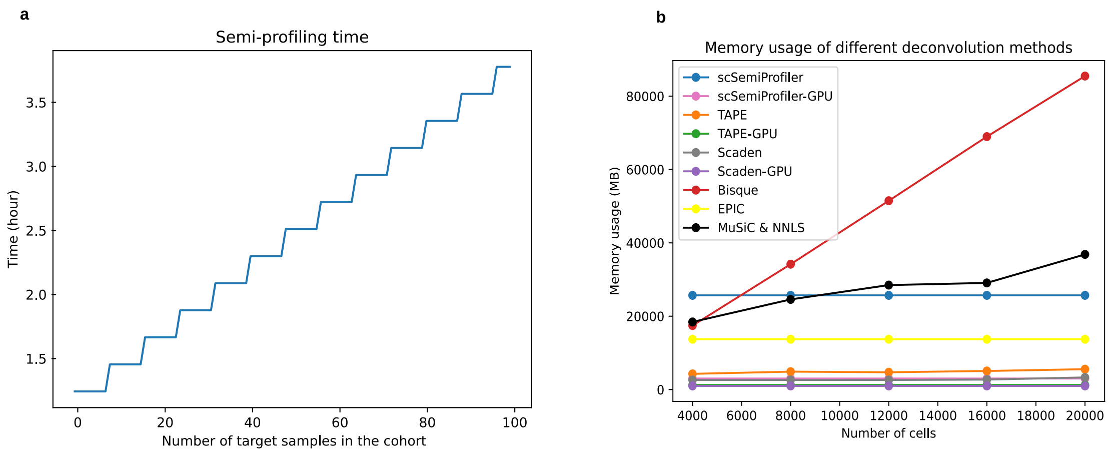


**Fig. S1**: **Runtime and memory usage.** The statistics are measured using our Linux server with 8 GPUs (4 Tesla M10 and 4 NVIDIA A16) and two Intel Xeon Platinum 8160 CPUs (each featuring 24 cores and supporting 2 threads per core, resulting in a total of 96 logical CPUs available for computation. **a**, Estimated runtime of scSemiProfiler when 4 representatives are sequenced and all 8 GPUs are used. The x-axis represents the number of target samples in the cohort, and the y-axis indicates time in hours. **b**, Comparison of memory usage among various deconvolution methods. Memory usage was monitored across deconvolution methods when using 1 to 5 samples (each has 4000 cells) as single-cell references for deconvoluting 4 bulk samples. The cell data matrix is sampled from the COVID-19 cohort and contains 6030 gene features. Most deep learning methods have very low memory usage and only increase slightly for reading larger input data matrices. Our method's memory usage does not increase with the number of samples, as different samples are trained using separate models sequentially. Bisque and MuSiC & NNLS exhibit nearly linear memory usage. CIBERSORTx, not being open-source software, offers its service exclusively through a website. Therefore, testing the exact memory usage is not feasible. The website permits a maximum upload of 1GB data per user, approximately equivalent to 42826 cells' gene expression data stored in TXT format, as required by the website.

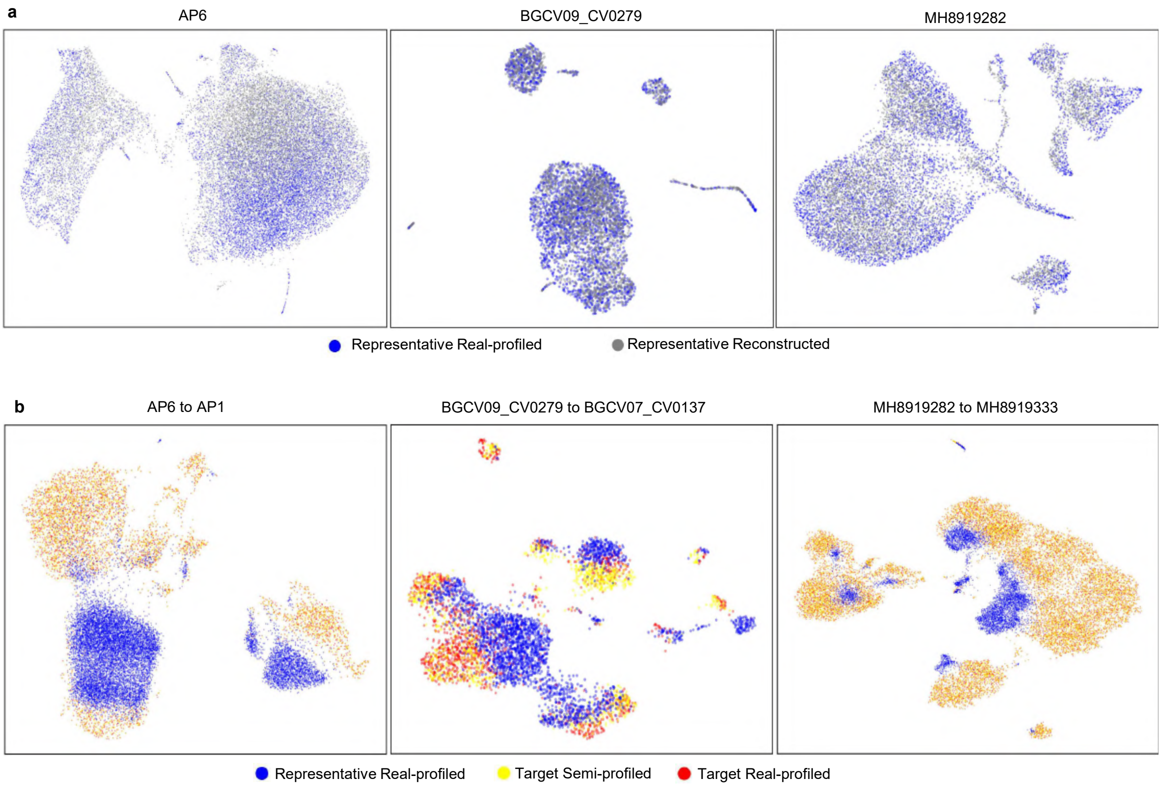


**Fig. S2**: **Examples of the in silico single-cell data inference for target samples in the COVID-19 cohort.** The first row shows the reconstruction of the single-cell data for 3 example representatives. The UMAP plots in the second row show the cell distribution of the representative, inferred target sample, and target sample ground truth single-cell data.

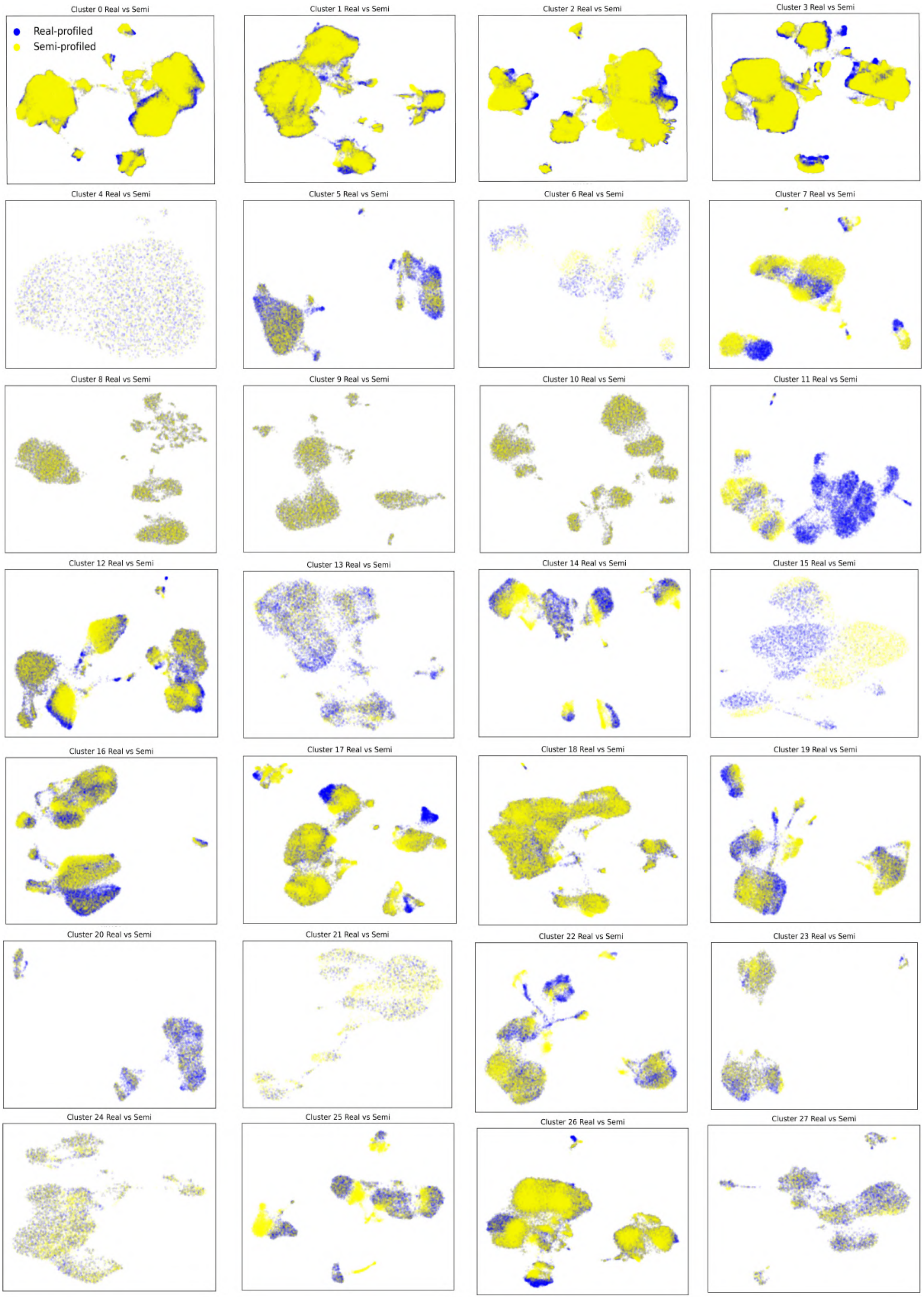


**Fig. S3**: **Comparison between the real-profiled COVID-19 dataset (blue) and the semi-profiled dataset (yellow) separated by sample clusters.** Each sample cluster includes a representative sample. Overlap area means the two versions of datasets are similar.

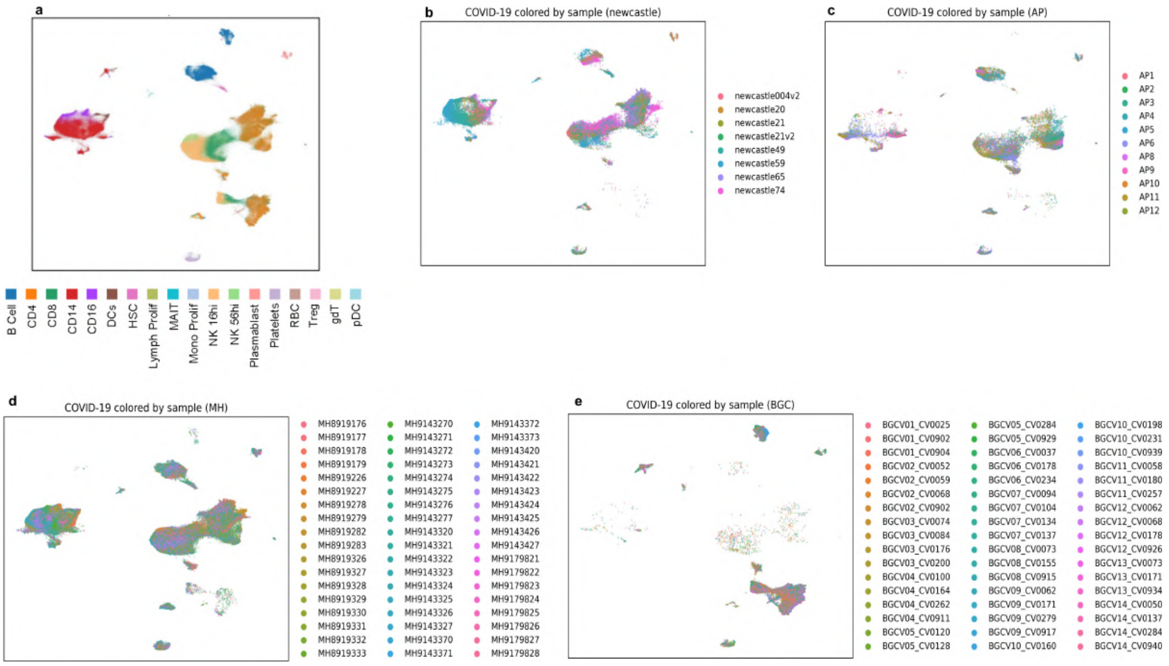


**Fig. S4**: **UMAP visualizations showing the cell distribution difference between different samples in the COVID-19 dataset. a**, UMAP of the real-profiled COVID-19 dataset colored by cell types. **b-e**, UMAP visualization of the real-profiled COVID-19 dataset colored by samples. By comparing with (**a**), it can be observed that the cell type distributions in samples are different.

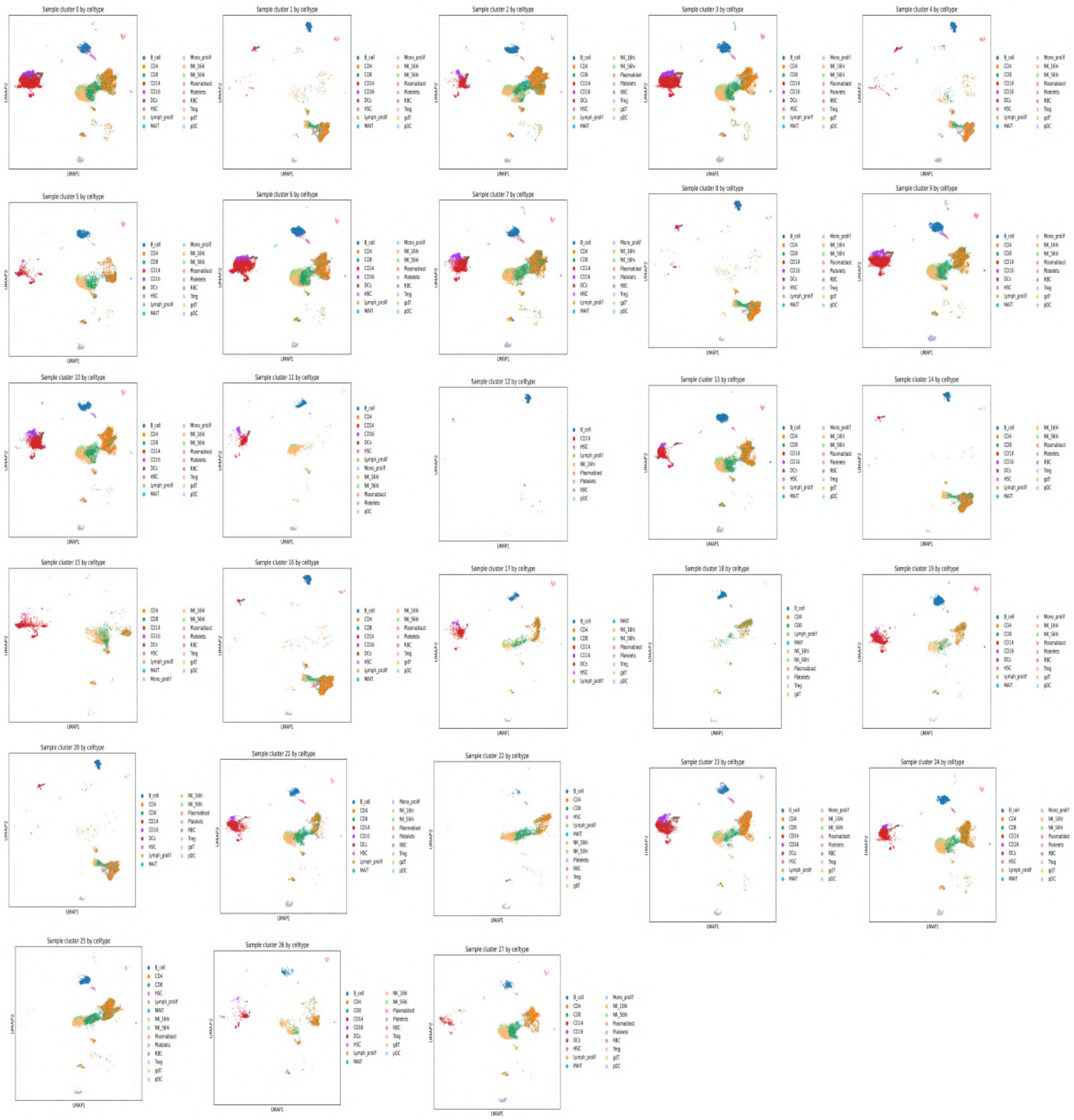

**Fig. S5**: **UMAP visualization of each sample cluster in the COVID-19 dataset, colored by cell types.** Cell type distributions vary substantially among different sample clusters.

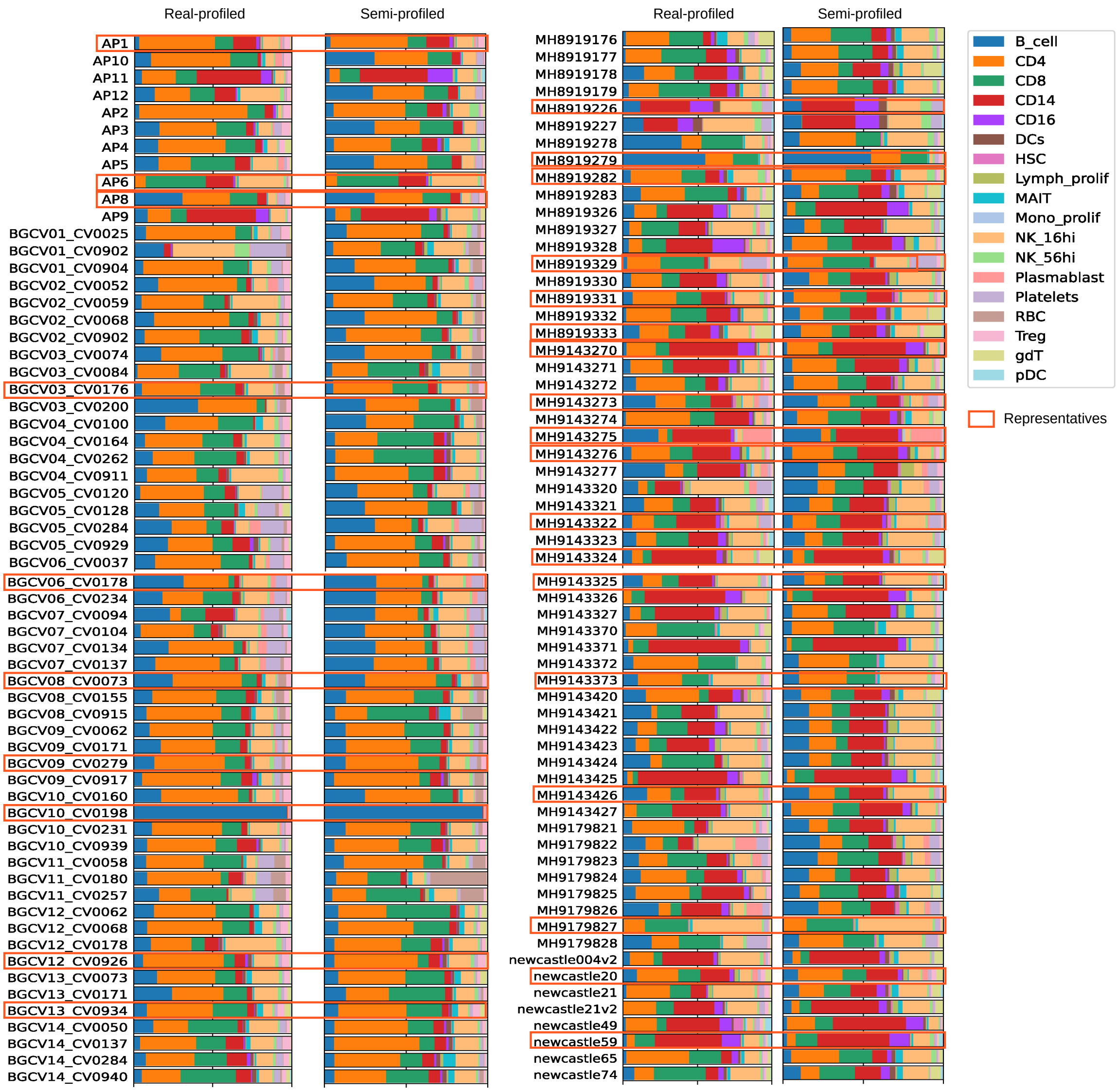


**Fig. S6**: **Deconvolution results for all samples in the COVID-19 cohort.** The average Pearson Correlation between the real-profiled and semi-profiled versions is 0.908, and the RMSE (Root Mean Square Error) is 0.146.

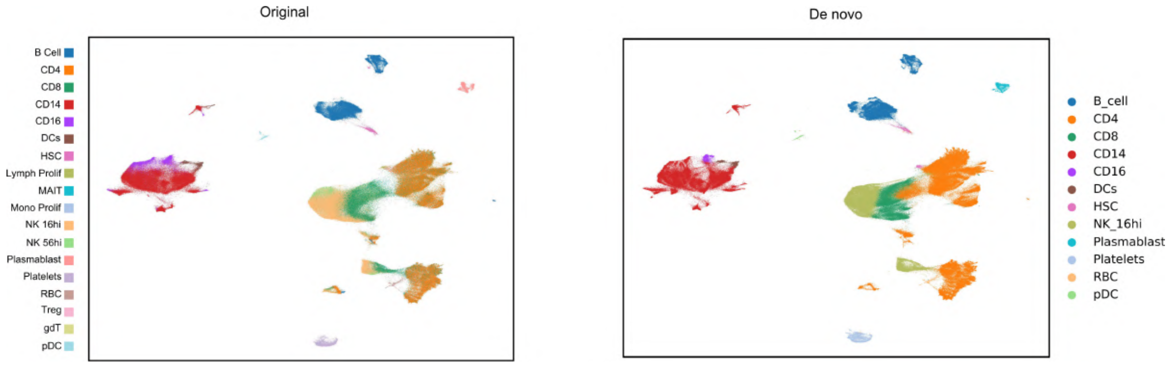


**Fig. S7**: **Side-by-side comparison of the real-profiled data and semi-profiled data annotated by *de novo* annotation.** The semi-profiled cohort annotated via supervised cell type annotation can be found in Fig. 2**b**.

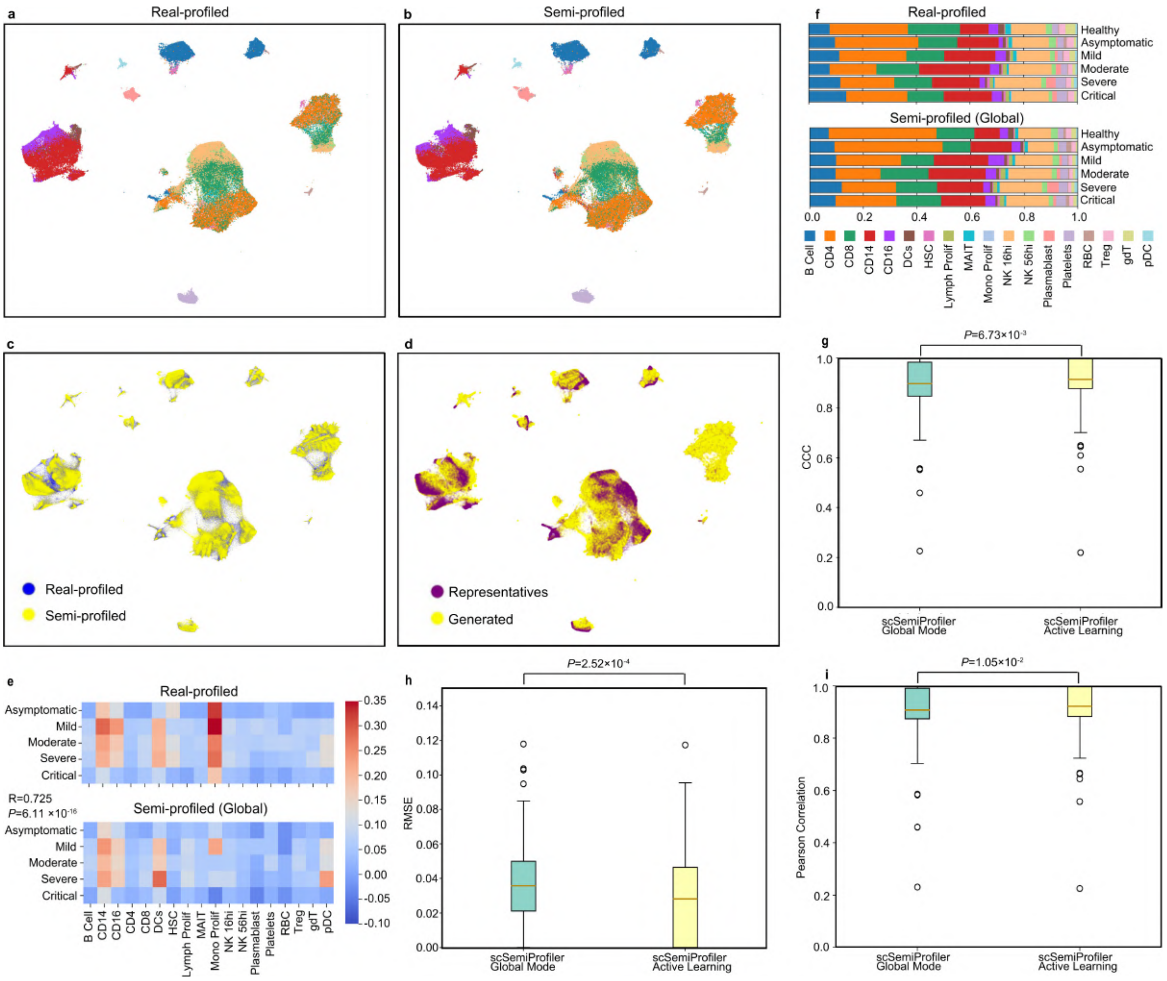


**Fig. S8**: **Comparative analyses between real-profiled COVID-19 datasets and the semi-profiled dataset under global mode. a**, UMAP visualization of the real-profiled COVID-19 cohort. Colors represent cell types and are consistent with (**f**). **b**, UMAP visualization of the global mode semi-profiled COVID-19 cohort. **c**, Combined UMAP visualization comparing the cell distribution across both datasets. **d**, UMAP visualization of the global mode semi-profiled dataset with different colors representing the representatives' cells selected by the global selection and the *in silico* inferred cells. **e**, Comparison of the interferon pathway activation pattern in both datasets. **f**, Cell type composition visualization in different COVID-19 disease severity levels. The global mode semi-profiled dataset shows similar cell type proportions with the real-profiled version in each severity level. The Pearson correlations are: Healthy: 0.959, Asymptomatic: 0.968, Mild: 0.991, Moderate: 0.985, Severe: 0.996, Critical: 0.979. **g-i**, Deconvolution performance comparison between semi-profiling using active learning and semi-profiling using global mode.

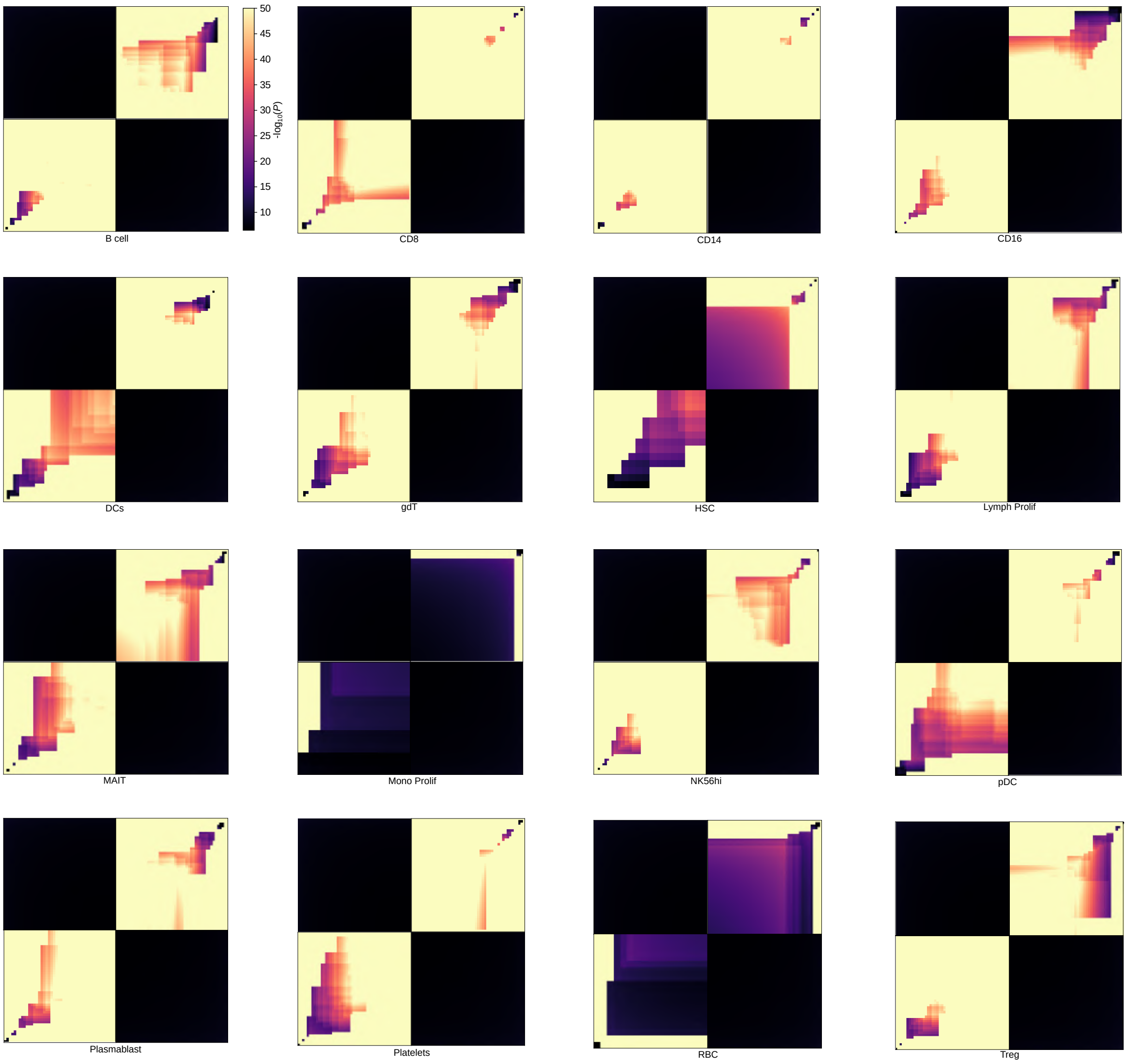


**Fig. S9**: **The Rank–rank hypergeometric overlap (RRHO) plots for other cell types in the COVID-19 dataset.** An RRHO plot visualizes the overlap between two ranked gene lists, highlighting the degree of similarity and the significance of the overlap between them. See more details about the RRHO plot in the Methods section. The top 50 positive and negative markers, identified using real-profiled and semi-profiled datasets, are utilized for the plots.

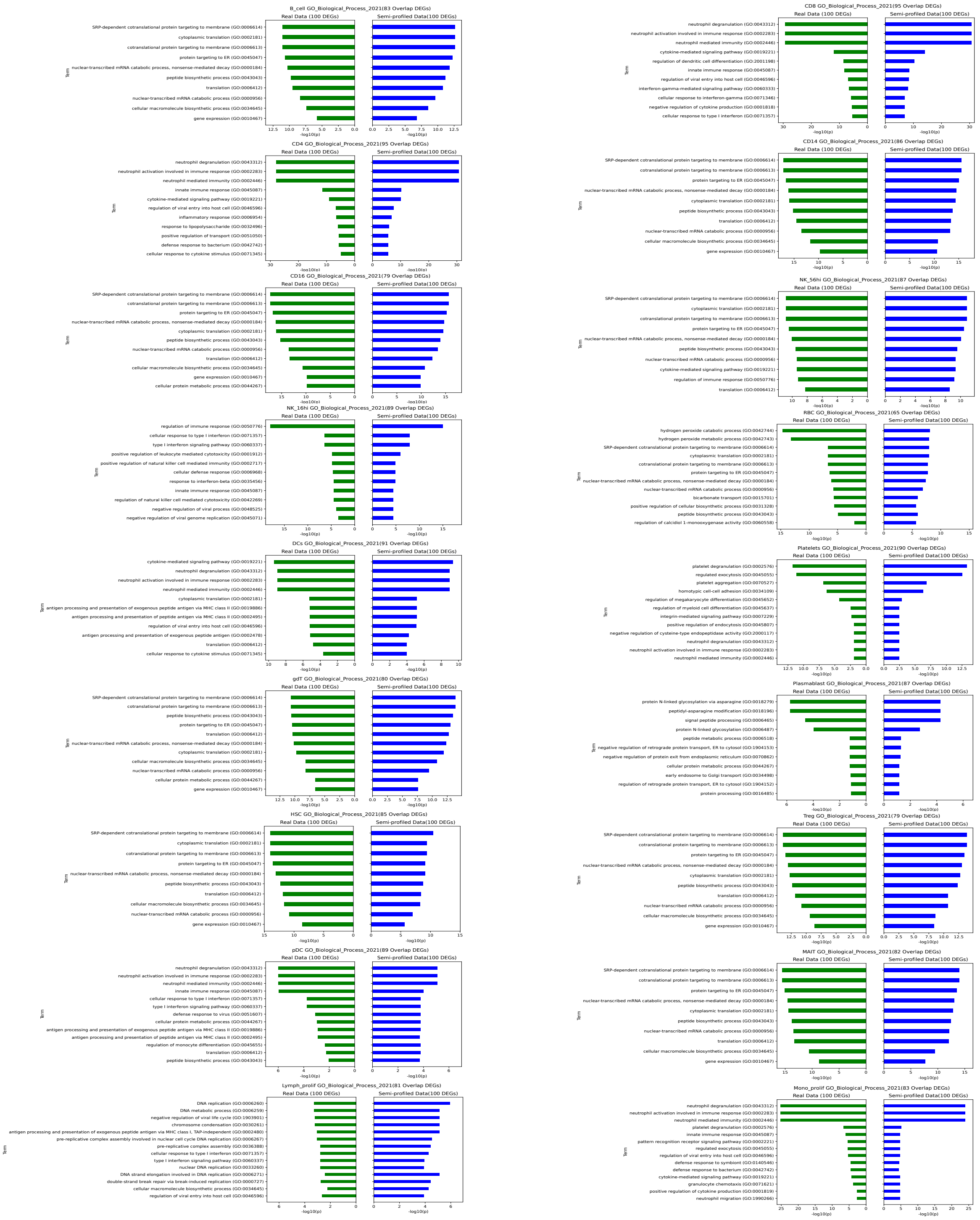

**Fig. S10**: **GO enrichment analysis for additional cell types in the COVID-19 dataset.** The top 100 signature genes are used for the analysis. The Pearson Correlations between the negative logged p-values (bar lengths) in the real-profiled and semi-profiled versions for each cell type are as follows: B_cell: 0.999, CD8: 0.999, CD14: 0.980, CD16: 0.994, DCs: 0.987, gdT: 0.985, HSC: 0.964, Lymph_prolif: 0.592, MAIT: 0.999, Mono_prolif: 0.993, NK_16hi: 0.974, NK_56hi: 0.996, pDC: 0.839, Plasmablast: 0.982, Platelets: 0.986, RBC: 0.656, Treg: 0.984

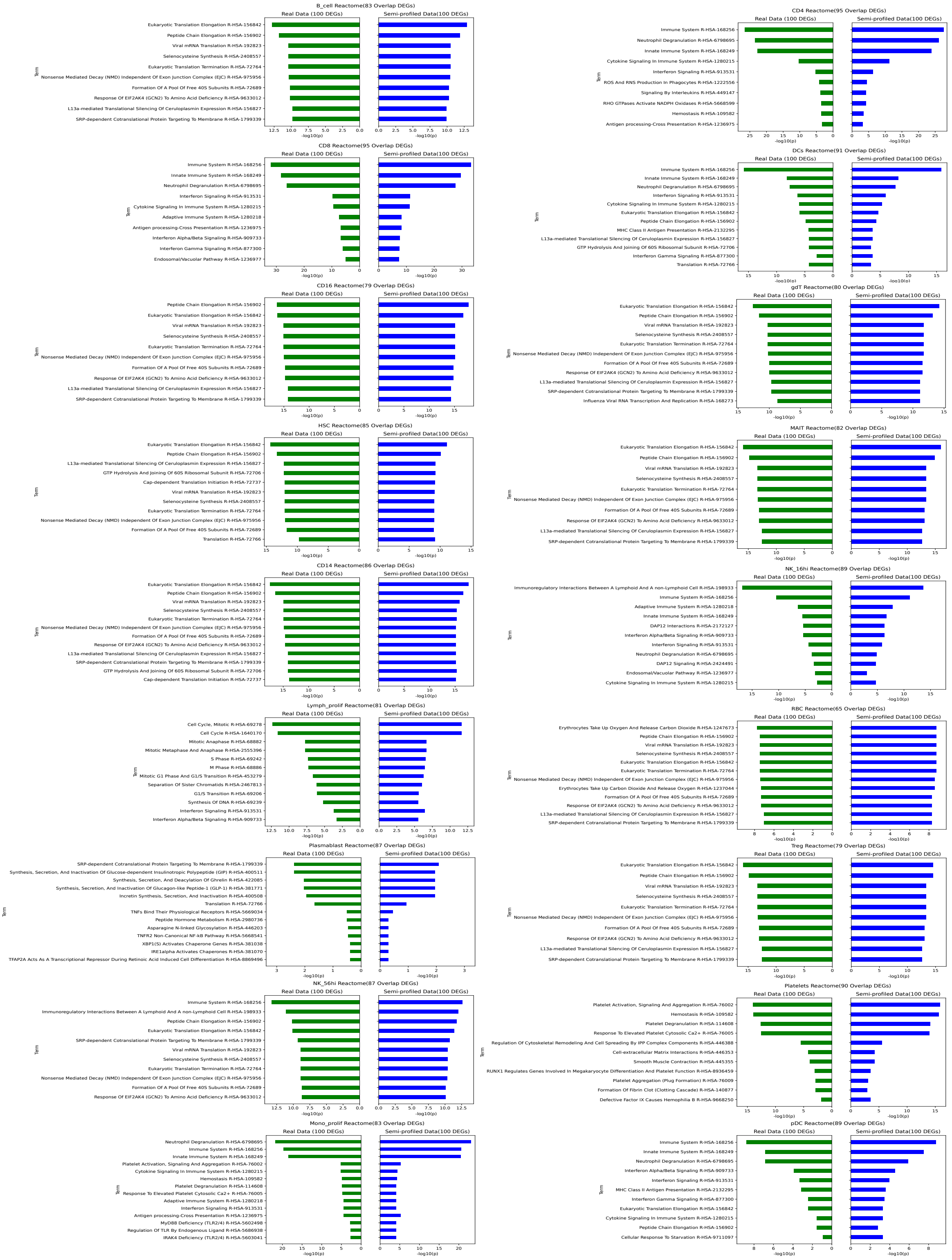


**Fig. S11**: **Reactome enrichment analysis for additional cell types in the COVID-19 dataset.** The top 100 signature genes are used for the analysis. The Pearson Correlations between the negative logged p-values (bar lengths) in the real-profiled and semi-profiled versions for each cell type are as follows: B_cell: 0.999, CD4:0.999, CD8: 0.999, CD14: 0.942, CD16: 0.963, DCs: 0.989, gdT: 0.961, HSC: 0.754, Lymph_prolif: 0.891, MAIT: 0.999, Mono_prolif: 0.993, NK_16hi: 0.968, NK_56hi: 0.952, pDC: 0.962, Plasmablast: 0.974, Platelets: 0.996, RBC: 0.767, Treg: 0.961

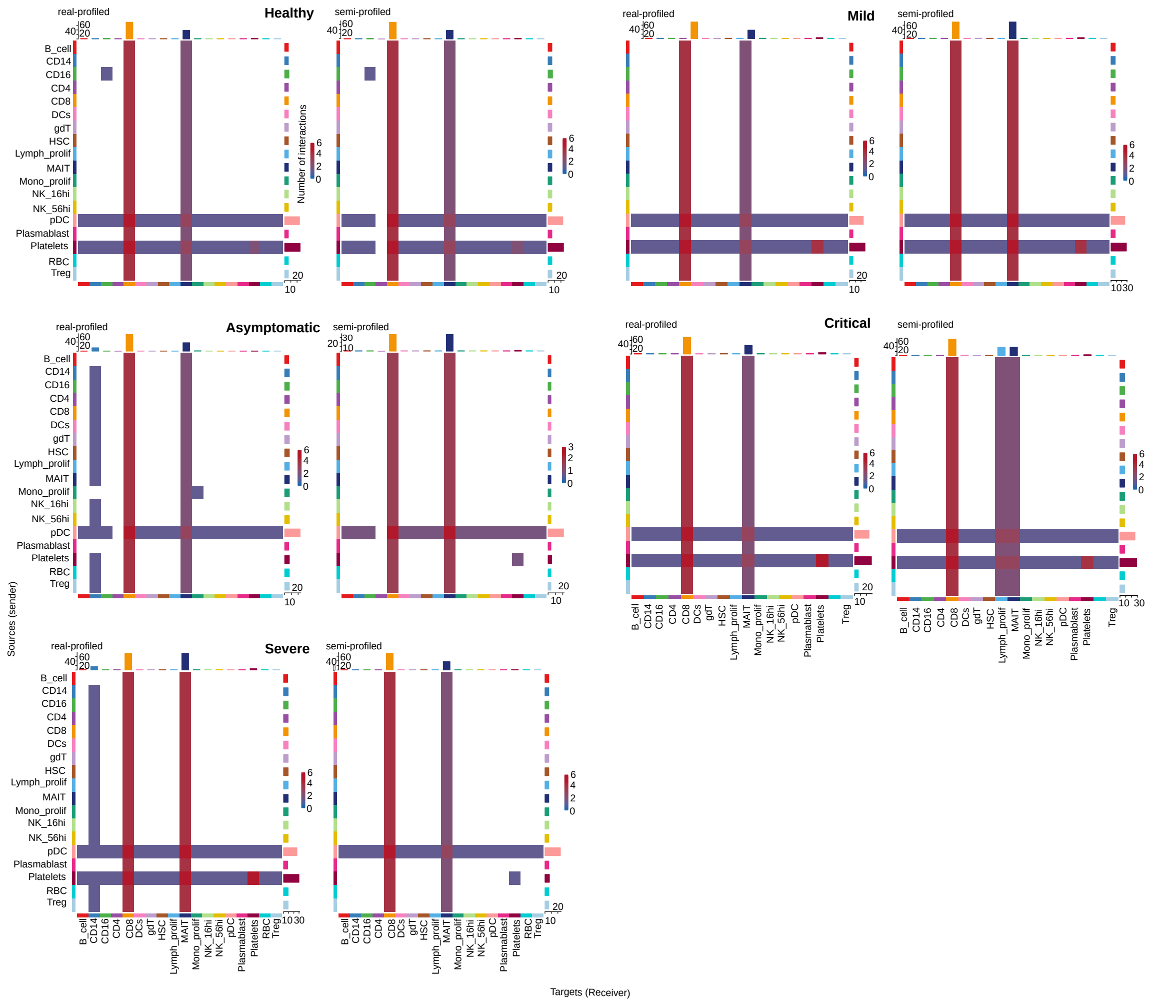


**Fig. S12**: **Comparison of cell-cell interaction analysis results across all severity levels in the COVID-19 dataset.** Pearson Correlations between the real and semi-profiled versions for each severity level are as follows: Healthy: 0.997, Asymptomatic: 0.915, Mild: 0.946, Moderate: 0.996, Severe: 0.908, Critical: 0.913.

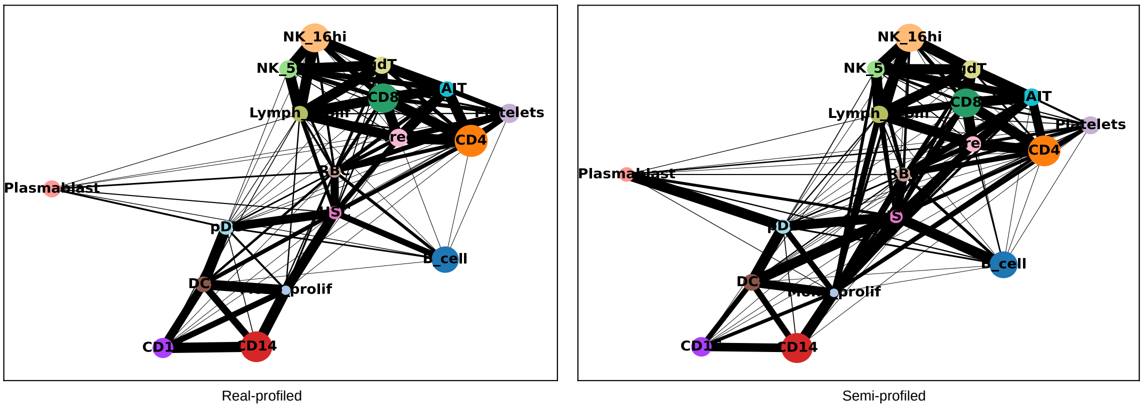


**Fig. S13**: **Comparison of PAGA results for the COVID-19 dataset.** The Pearson Correlation between the connectivity matrices of the PAGA plots derived from the real and semi-profiled datasets is 0.893. For easier comparison, the positions of the nodes are synchronized between the ground truth and the semi-profiled PAGA results.

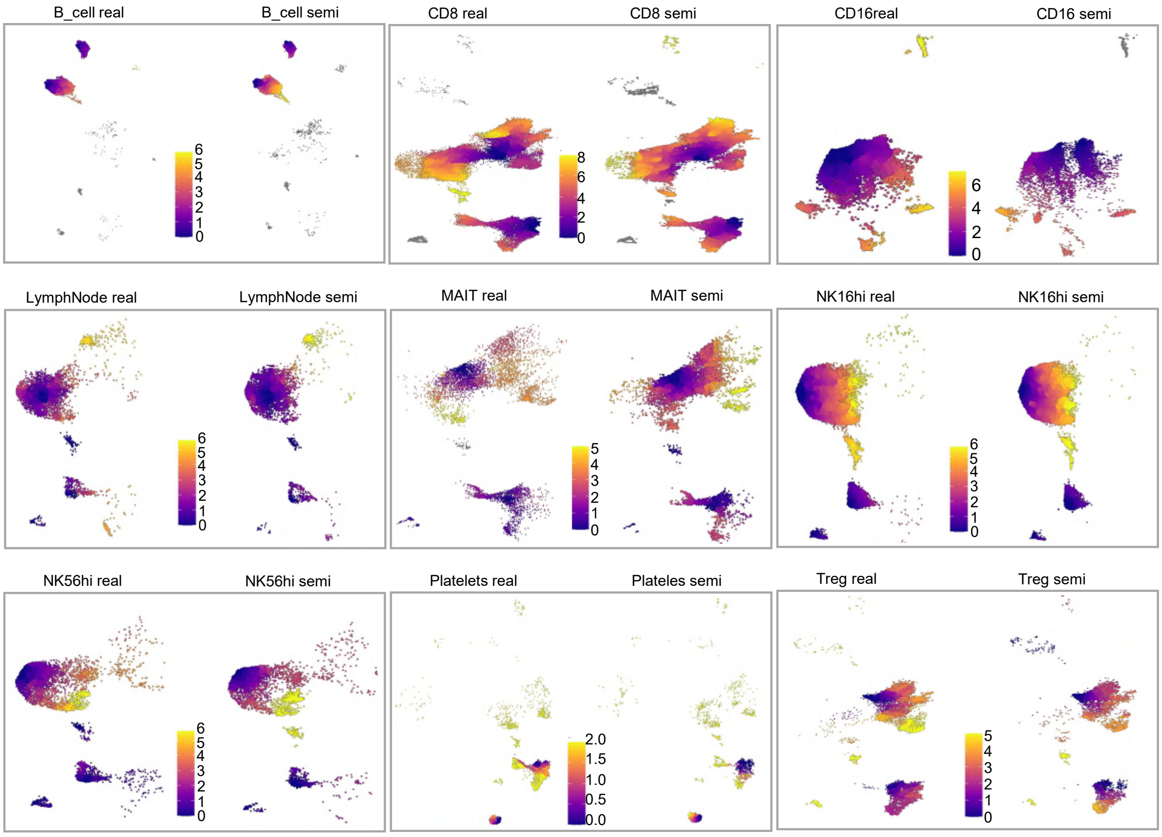


**Fig. S14**: **Pseudotime analysis results comparison for other cell types in the COVID-19 dataset.** Cell types with insufficient cells to form a cluster structure were excluded.

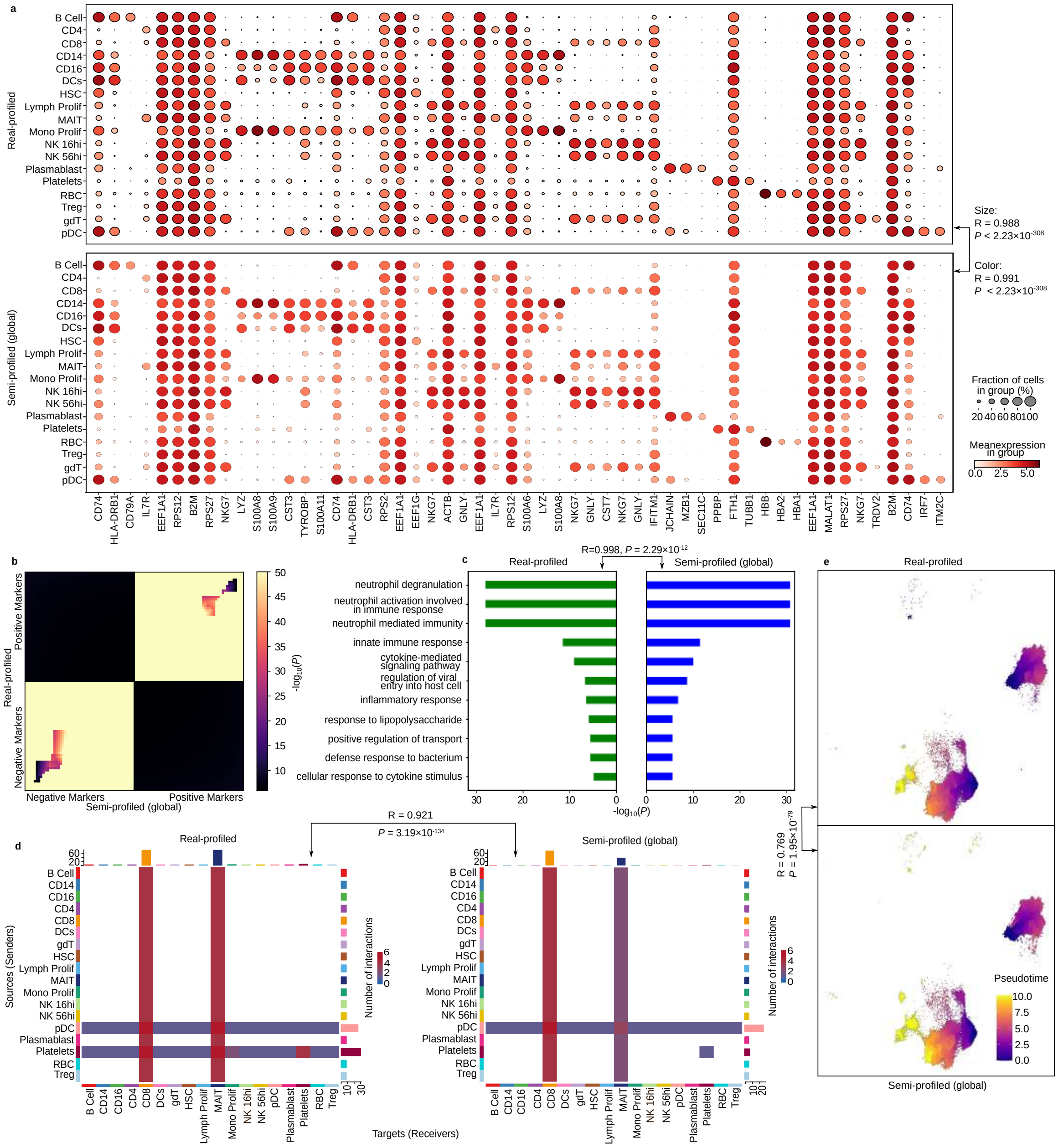


**Fig. S15**: **Comparative analyses of single-cell level downstream analysis tasks using the real-profiled COVID-19 dataset and semi-profiled COVID-19 dataset generated using the global mode.** **a**, Dot plots visualizing the top cell type signature genes' expression in real-profiled data (top) and global mode semi-profiled data (bottom). **b**, RRHO plot comparing the CD4 positive and negative markers in both datasets. **c**, CD4 cell type signature genes GO enrichment analysis results comparison. **d**, Cell-cell interaction analysis results comparison between the real-profiled dataset and the global mode semi-profiled dataset. **e**, Pseudotime results comparison using the real-profiled and global mode semi-profiled CD4 cells.

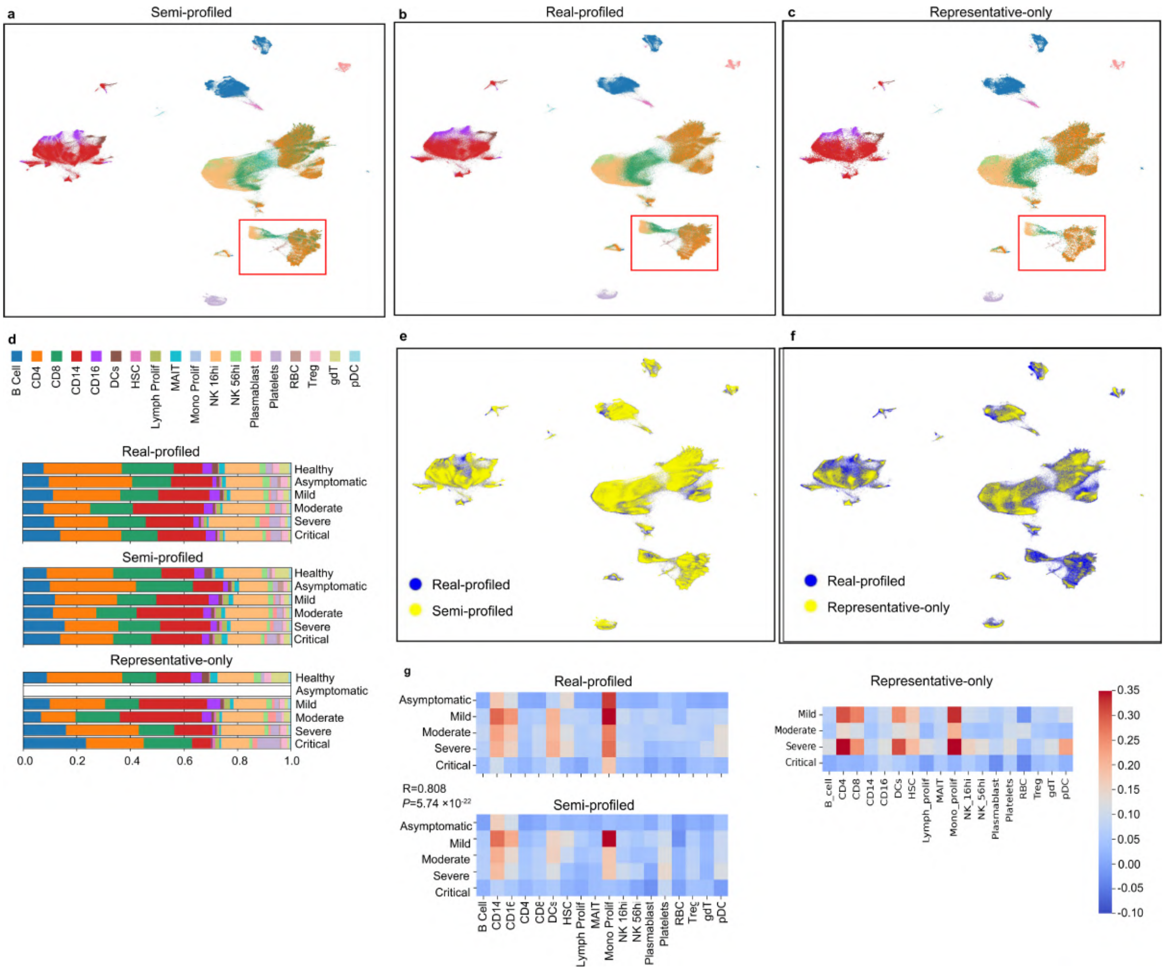


**Fig. S16**: **Comparing the semi-profiled COVID-19 cohort, real-profiled cohort, and representative-only cohort. a-c**, UMAP visualization of the semi-profiled cohort, real-profiled cohort, and representative-only cohort (only representative cells are included) respectively. Different colors represent different cell types and are consistent with (**d**). **d**, Stacked bar plots visualizing different cohorts' cell type proportion in different COVID-19 severity level. The semi-profiled cohort shows high similarity with the real-profiled cohort, the Pearson correlations between the semi-profiled cohort and real-profiled cohort in each condition are: Healthy: 0.987, Asymptomatic: 0.970, Mild: 0.996, Moderate: 0.992, Severe: 0.978, Critical: 0.989. Representative-only cohort shows a lower similarity with the real-profiled cohort. Pearson correlations: Healthy: 0.971, Asymptomatic: N/A, Mild: 0.965, Moderate: 0.981, Severe: 0.961, Critical: 0.859. **e**, Joint UMAP showing the similarity between the semi-profiled cohort and the real-profiled cohort. **f**, Joint UMAP the representative-only cohort fails to cover some areas of the real-profiled cohort's UMAP. **g**, Interferon pathway activation pattern comparison between the three cohorts. The representative-only cohort cannot generate results for "Asymptomatic".

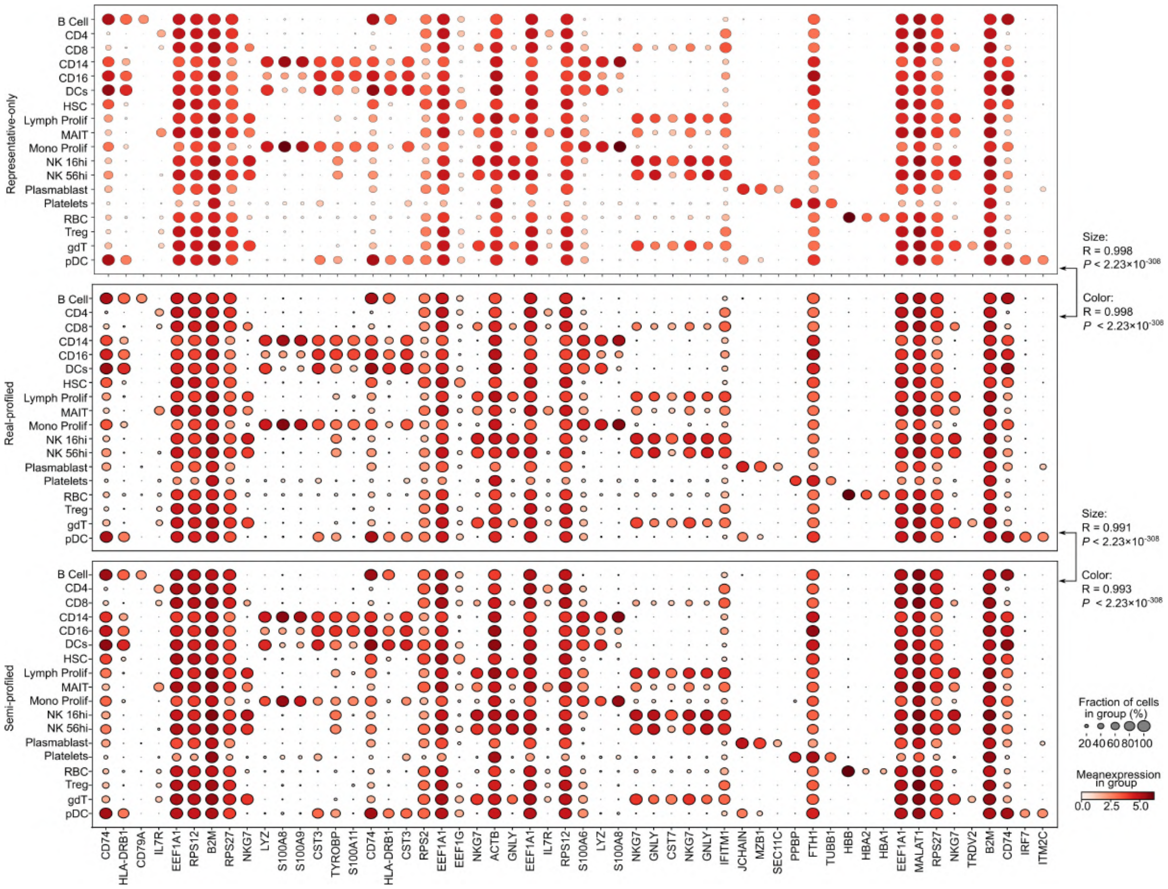


**Fig. S17**: **Dot plots visualizing the top cell type signature markers' expression patterns in the semi-profiled COVID-19 cohort, real-profiled cohort, and the representative-only cohort.**

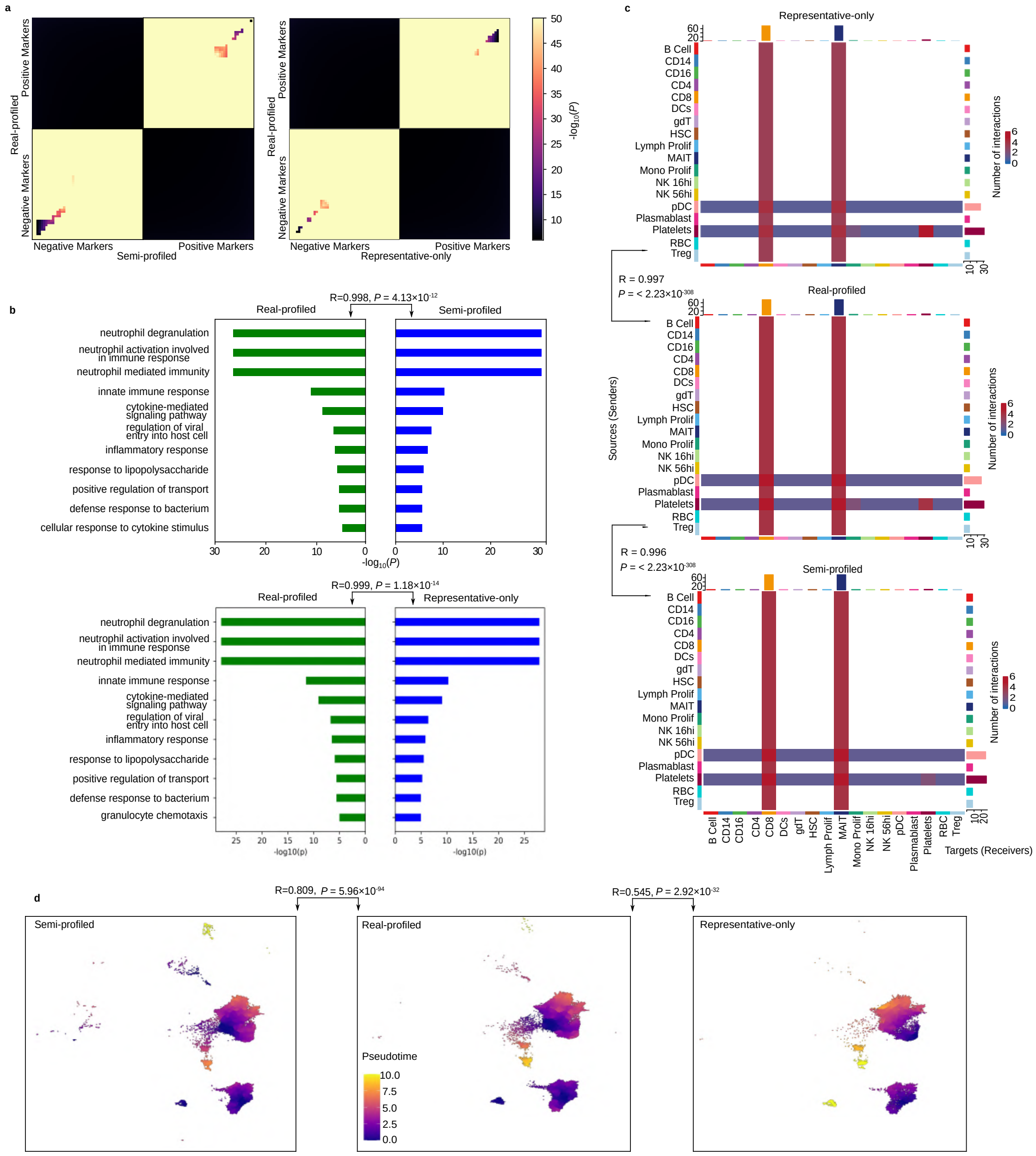


**Fig. S18**: **Downstream single-cell analysis results using real-profiled COVID-19 cohort, semi-profiled cohort, and representative-only cohort. a**, RRHO plots comparing the similarity between CD4 markers found using real-profiled cohort and semi-profiled cohort (left) and the similarity between CD4 markers found using real-profiled cohort and representative-only cohort. **b**, Comparison of GO enrichment analysis using CD4 genes found by different cohorts (top: real-profiled vs. semi-profiled, bottom: real-profiled vs. representative-only). **c**, Comparing the cell-cell interaction analysis results between the three cohorts. **d**, Pseudotime analysis results between the real-profiled cohort and semi-profiled cohort are more similar (Pearson correlation: 0.809) than that between real-profiled cohort and representative-only cohort (Pearson correlation: 0.545).

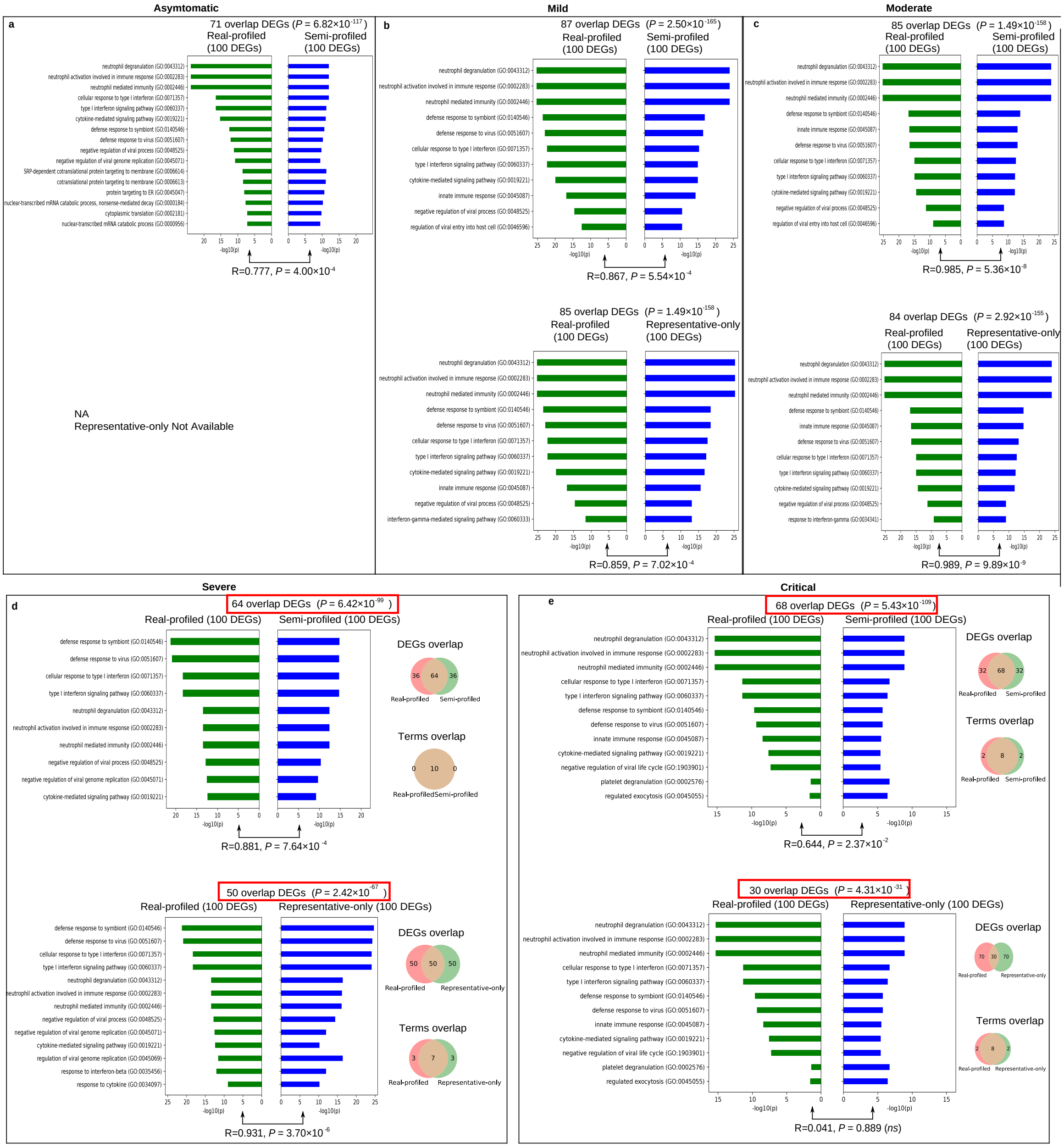


**Fig. S19**: **Evaluating the disease condition markers identified using the semi-profiled COVID-19 cohort and only the 28 representatives by comparing against the real-profiled cohort.** Markers are differentially expressed genes between the disease condition group and the healthy group in each cohort. a, Markers for "Asymptomatic" conditions identified in the semi-profiled cohort align closely with those from the real-profiled cohort; the representative-only cohort lacks data on "Asymptomatic" patients and hence did not perform this analysis. **b**, "Mild" markers comparison. **c**, "Moderate" markers comparison. **d-e**, Comparisons of "Severe" and "Critical" condition markers, respectively. Markers from the semi-profiled cohort demonstrate a significantly higher overlap with those from the real-profiled cohort compared to the representative-only cohort, reflecting greater similarity in the GO enrichment analysis results (more overlap in top 10 GO terms).

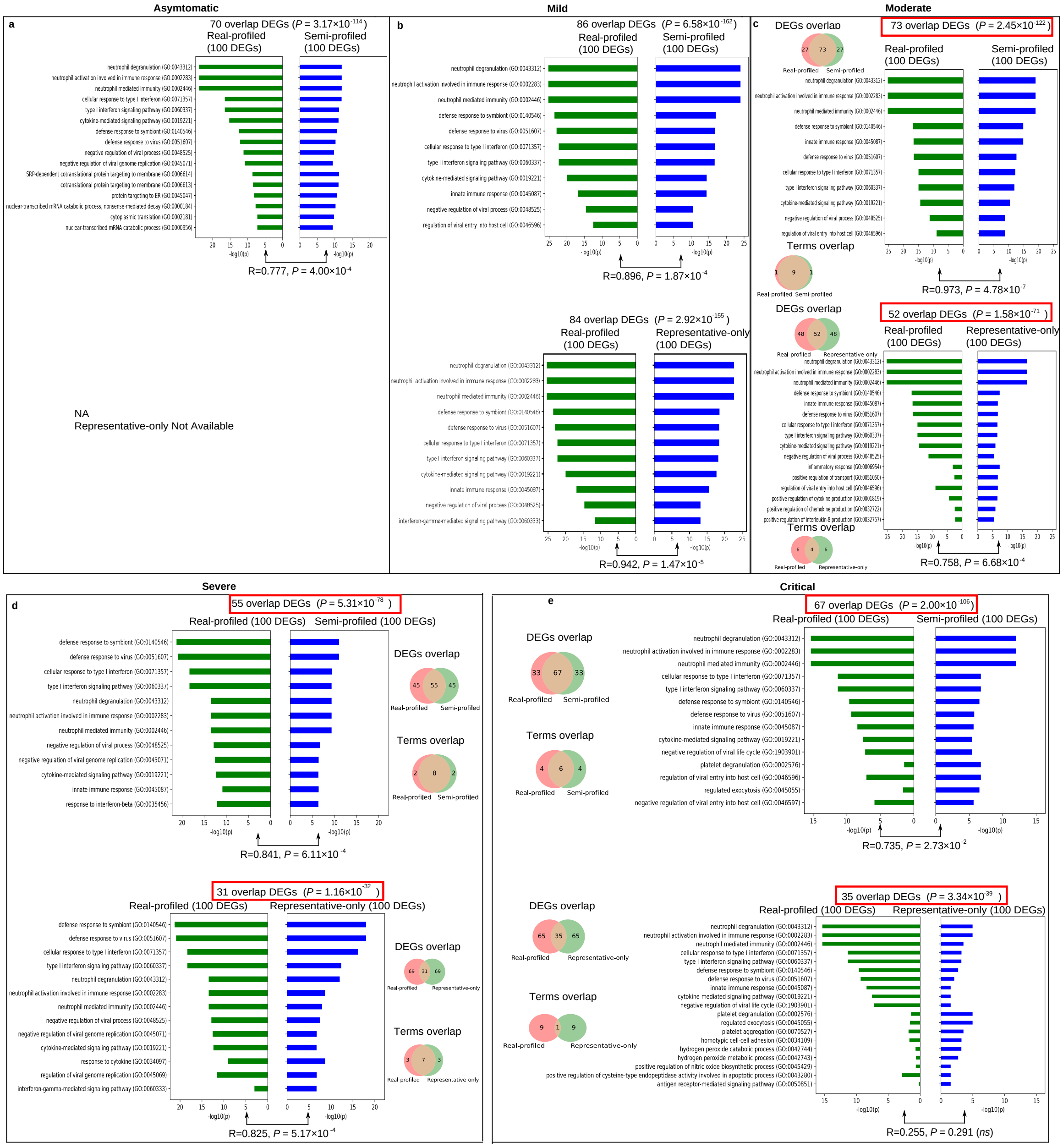


**Fig. S20**: **Comparing disease condition markers identified using three COVID-19 cohorts: the semi-profiled cohort based on 12 representatives, real-profiled cohort, and the only 12 representatives.** a, Despite the limited number of representatives, the semi-profiled cohort's "Asymptomatic" markers closely align with those from the real-profiled cohort. **b**, "Mild" markers comparison. **c-e**, Markers for "Moderate," "Severe," and "Critical" conditions in the semi-profiled cohort show significantly greater similarity to those in the real-profiled cohort, both in terms of marker overlap and GO enrichment analysis results compared to the representative-only cohort.

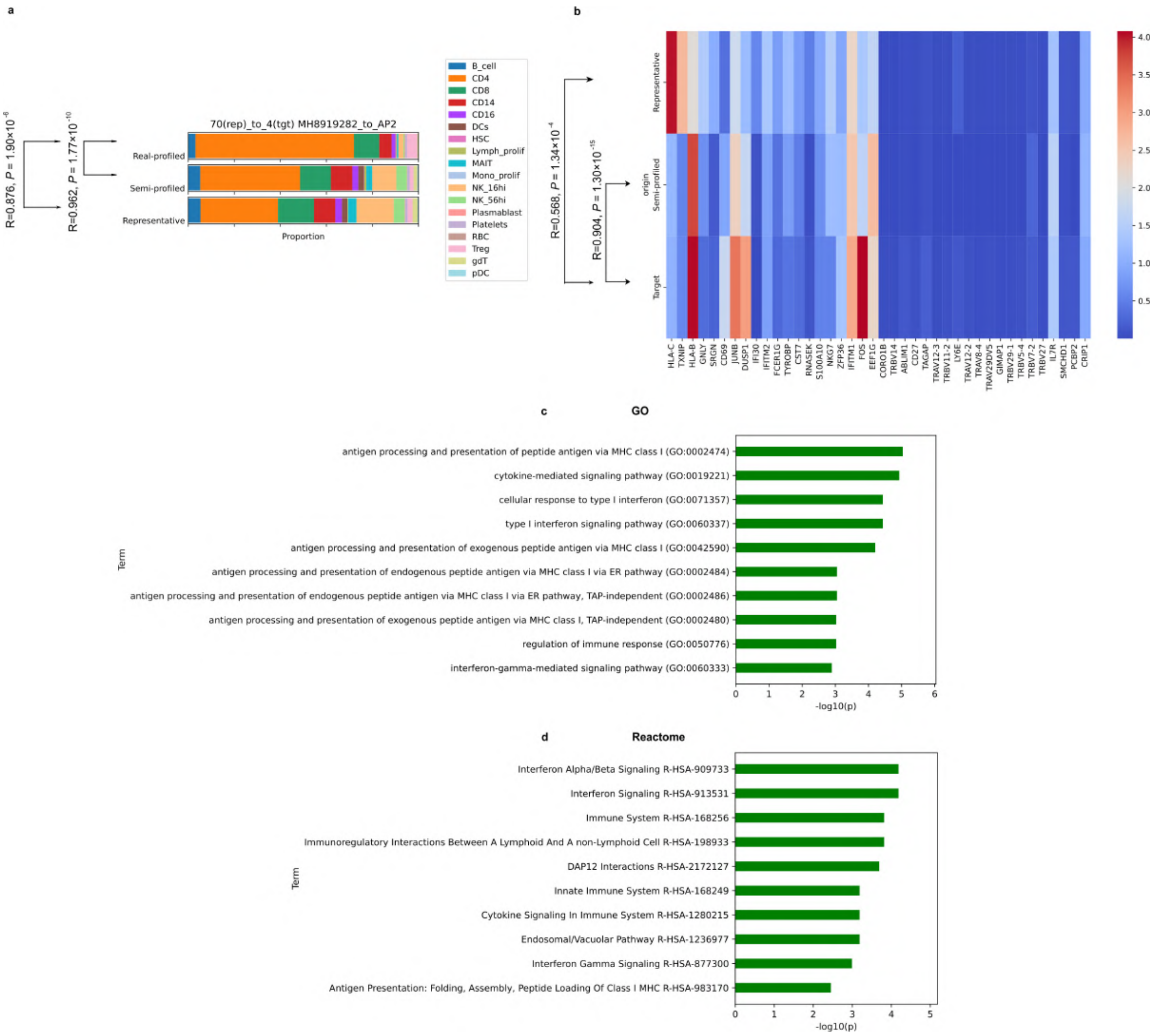


**Fig. S21**: **Semi-profiled cohort can provide a much more accurate representation of the ground truth cohort and reveal more biological insights as demonstrated in the COVID-19 dataset.** **a**, The inferred target sample with “Severe” COVID-19 in the semi-profiled cohort has significantly more similar cell type composition with the ground truth than the representative. **b**, The inferred target sample has a significantly more similar disease marker gene expression pattern with the ground truth than the representative. Markers are identified by performing differentially expressed gene analysis between the target ground truth sample and healthy controls. The left 20 are DEGs that show gene expression more similar to the target sample than the representatives, and the right 20 show where the representatives have more similar patterns than the inferred sample. **c-d**, GO and Reactome enrichment analysis using the left 20 DEGs in (**b**).

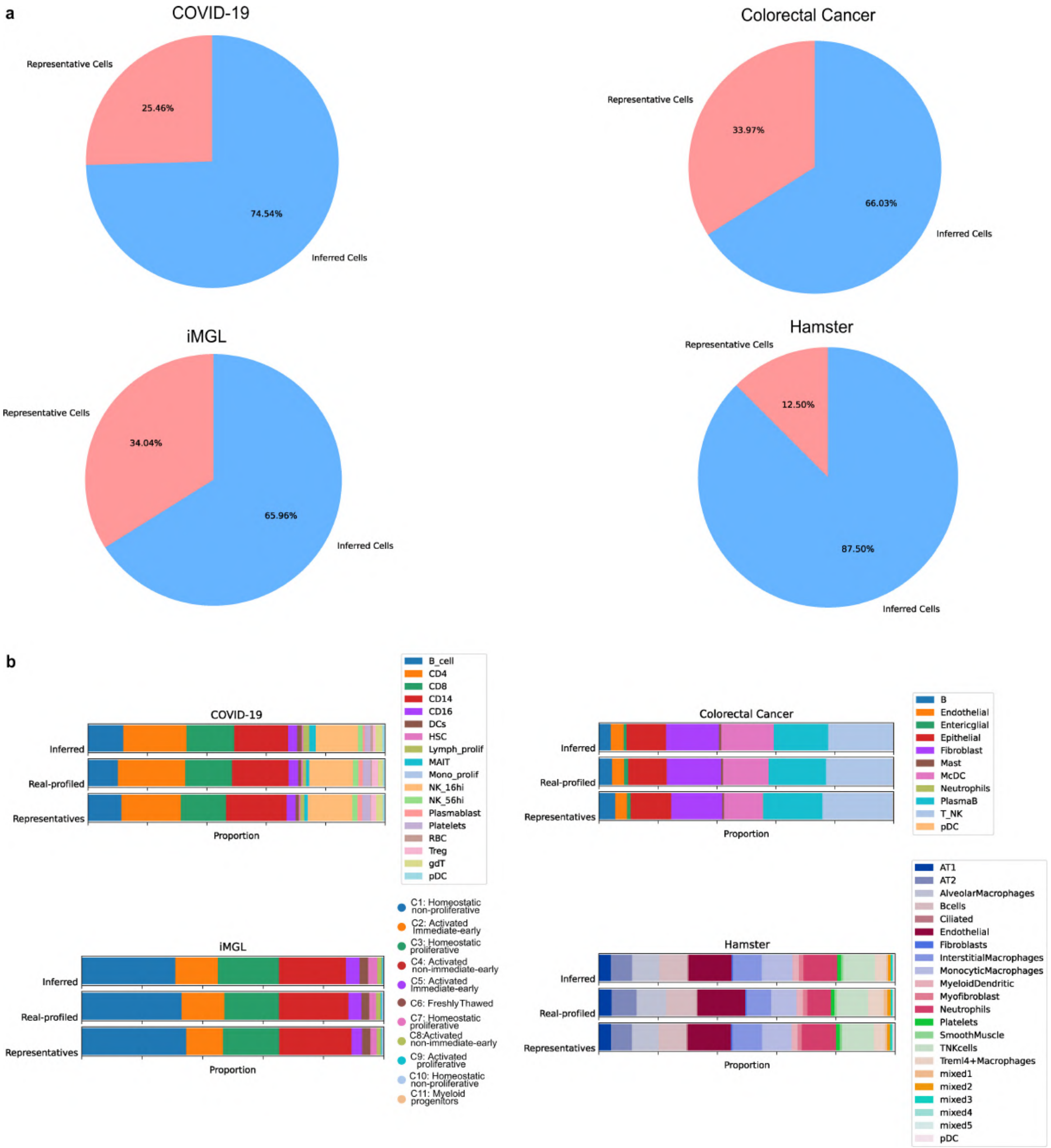


**Fig. S22**: **Overview of the real-profiled cells, representative cells, and *in silico* inferred cells in the four cohorts we analyzed. a**, Pie plots showing that in all semi-profiled cohorts, most cells are inferred using our deep generative model. **b**, Stacked bar plots showing the cell type proportions in the inferred-only cells, representative cells, and the whole real-profiled cohort are very similar.

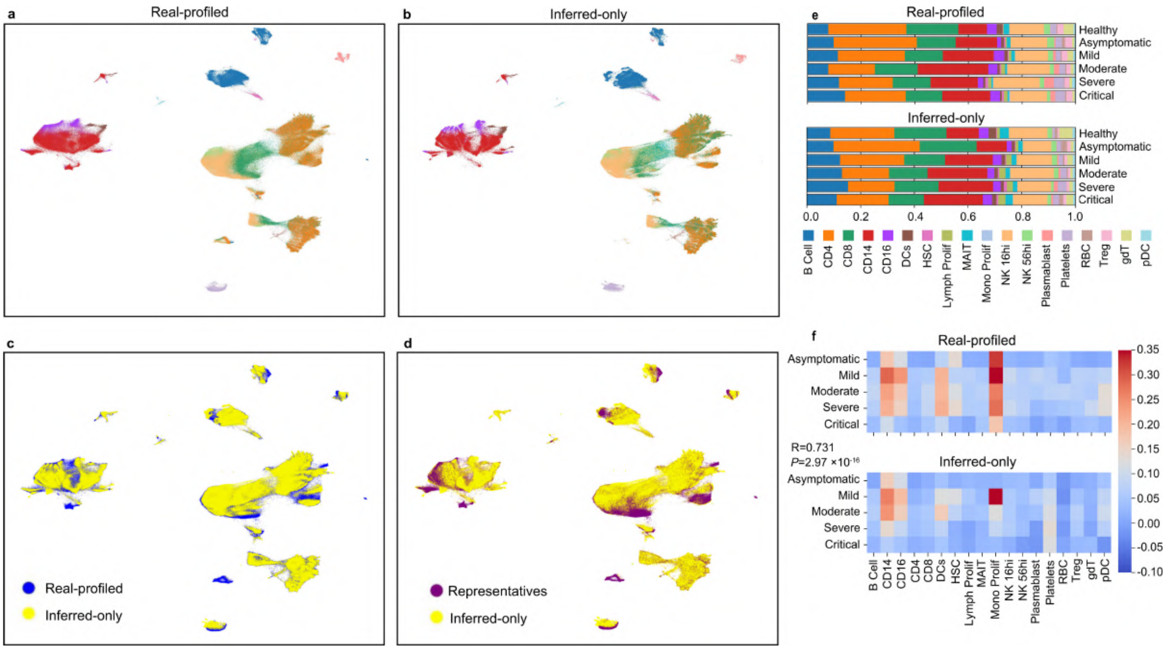


**Fig. S23**: **Similarity between the inferred-only dataset (representing the COVID-19 cohort using only *in silico* inferred cells) and the real-profiled dataset. a**, Real-profiled COVID-19 cohort UMAP visualization. Colors represent different cell types and are consistent with (**b**) and (**e**). **b**, UMAP visualization of the inferred-only dataset. **c**, UMAP visualization combining real-profiled cohort and inferred-only cells, showing the deep generative learning model is capable of generating cells similar to the real-profiled cohort for areas not covered well by the representatives. **d**, UMAP visualizing the representatives' cells and inferred-only cells. **e**, Cell type proportion comparison between the real-profiled dataset and the inferred-only dataset in different COVID-19 disease severity levels. The Pearson correlations are: Healthy: 0.982, Asymptomatic: 0.970, Mild: 0.995, Moderate: 0.975, Severe: 0.963, Critical: 0.970 **f**, Interferon pathway activation pattern comparison.

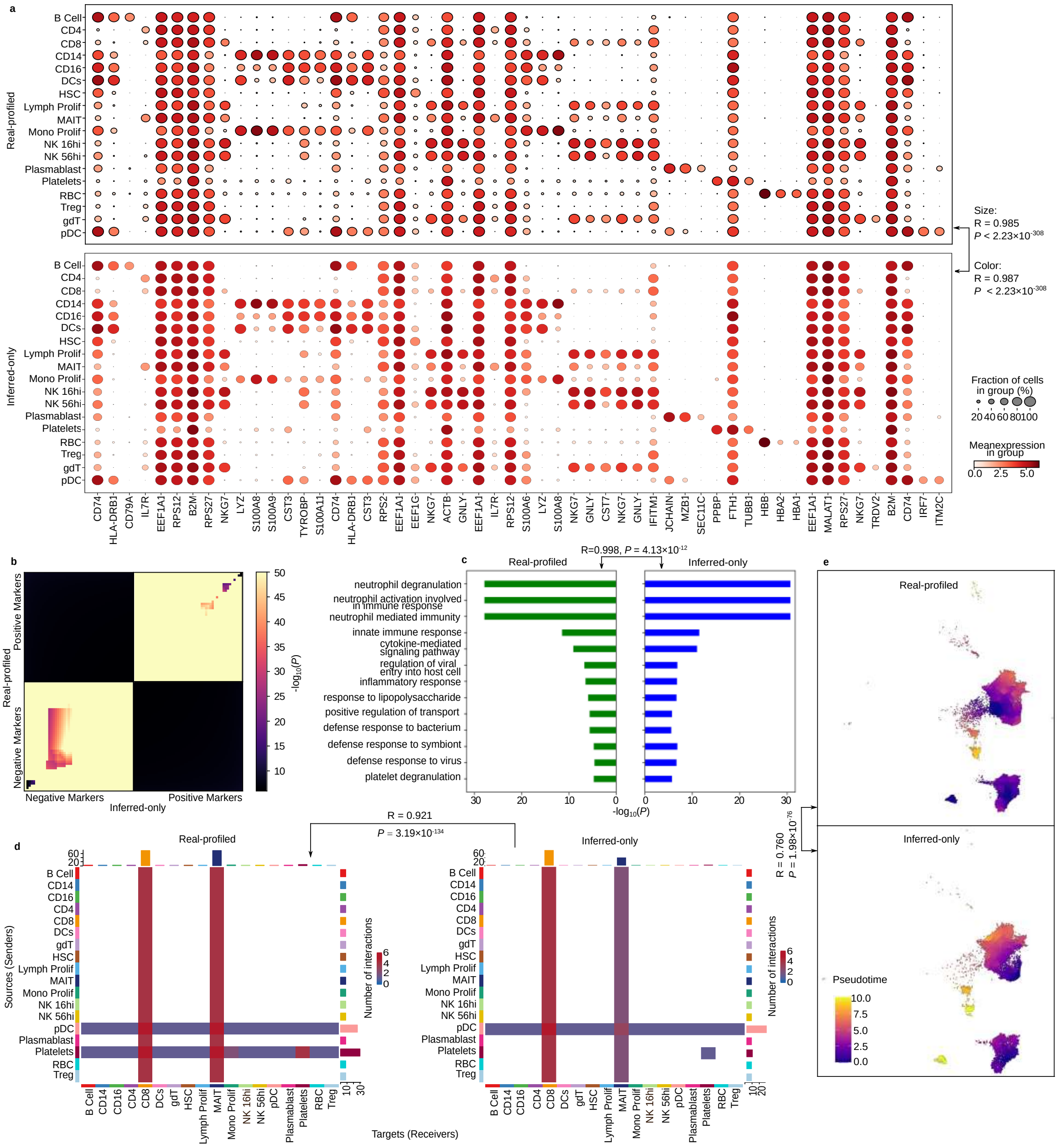

**Fig. S24**: **Downstream analysis using only *in silico* inferred cells (inferred-only dataset) generate similar results with the real-profiled COVID-19 cohort. a**, Dot plots showing nearly identical expression patterns of top cell type signature genes in the inferred -only dataset and real-profiled dataset. **b**, RRHO plot comparing the CD4 markers found using the two versions of datasets. **c**, CD4 cell type signature genes GO enrichment analysis results comparison. **d**, Cell-cell interaction analysis results comparison between the real-profiled dataset and inferred-only dataset. **e**, Pseudotime results comparison using the real-profiled and *in silico* generated CD4 cells.

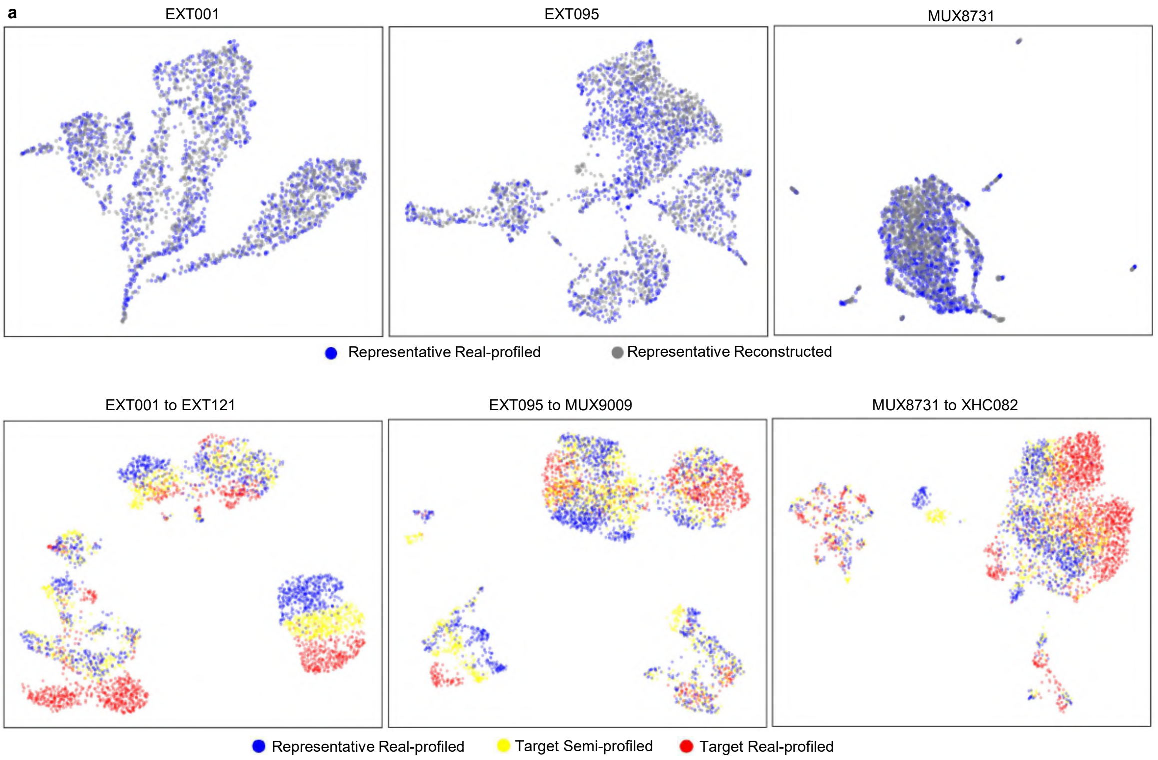


**Fig. S25**: **Examples of the in silico single-cell data inference for target samples in the colorectal cancer dataset.** The first row shows the reconstruction of the corresponding representatives. The second row shows the cell distribution of the representative, inferred target sample, and target sample ground truth.

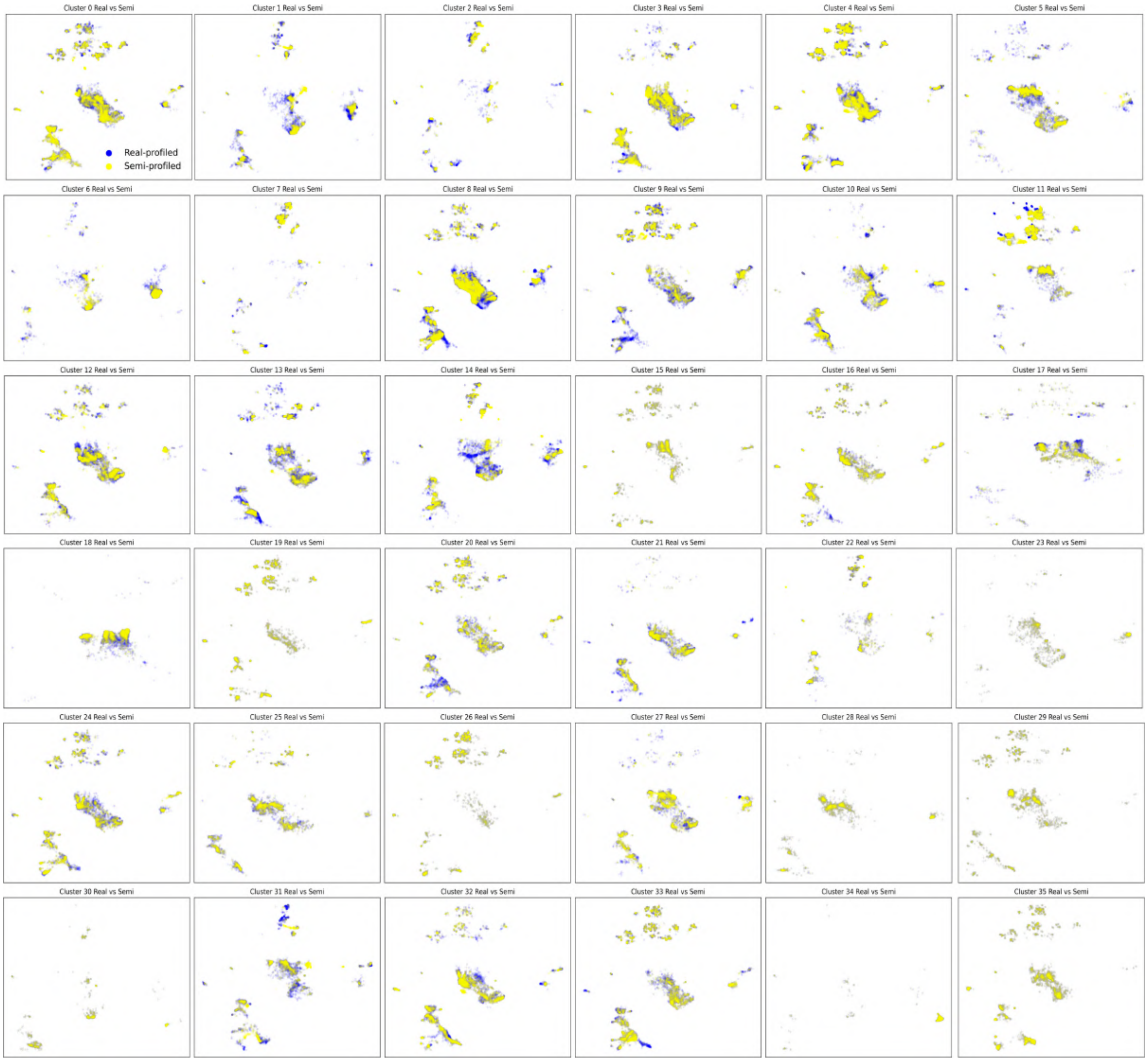


**Fig. S26**: **Comparison between the real-profiled colorectal cancer dataset (blue) and semi-profiled dataset (yellow) separated by sample clusters.**

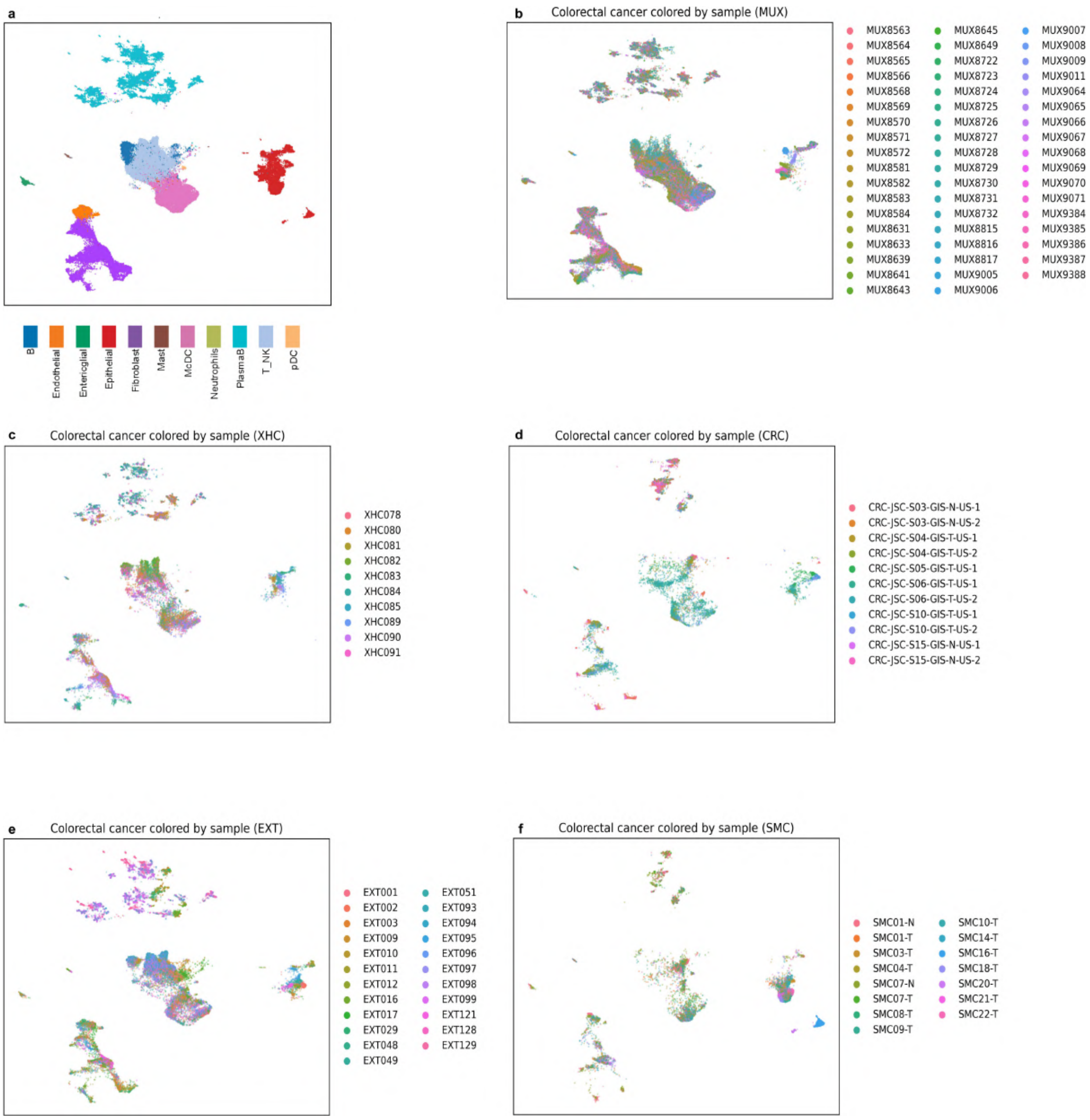


**Fig. S27**: **Visualizing cells from different samples in the colorectal cancer dataset. a**, UMAP visualization of the real-profiled colorectal cancer dataset colored by cell types. **b-f**, UMAP visualization of the real-profiled colorectal cancer dataset colored by samples. Samples from different batches are plotted separately. By comparing with (**a**), it can be observed that the cell type distributions in samples are different.

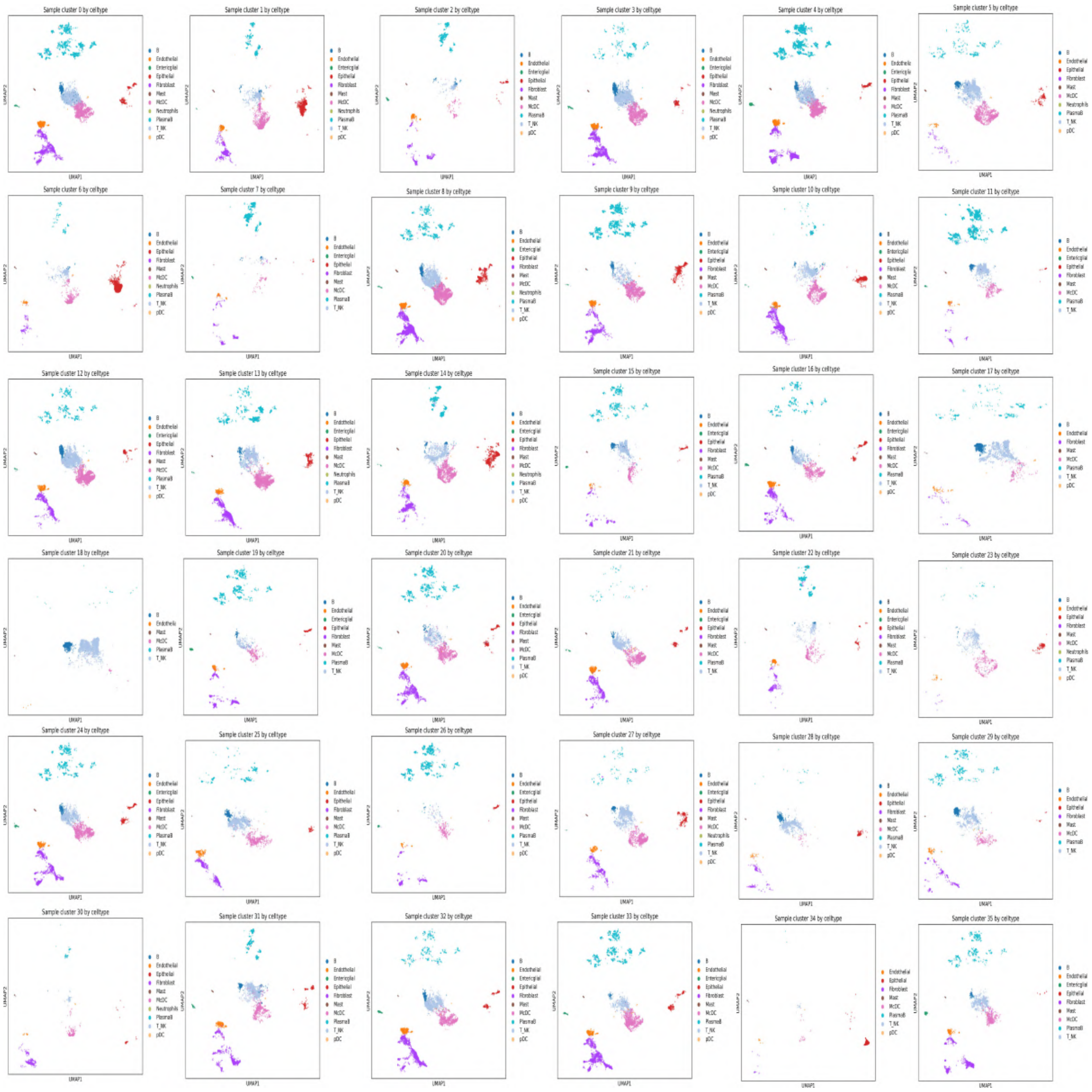

**Fig. S28**: **UMAP visualization of each sample cluster in the colorectal cancer dataset.** Colors represent cell types. The distributions of cell types are different in different sample clusters.

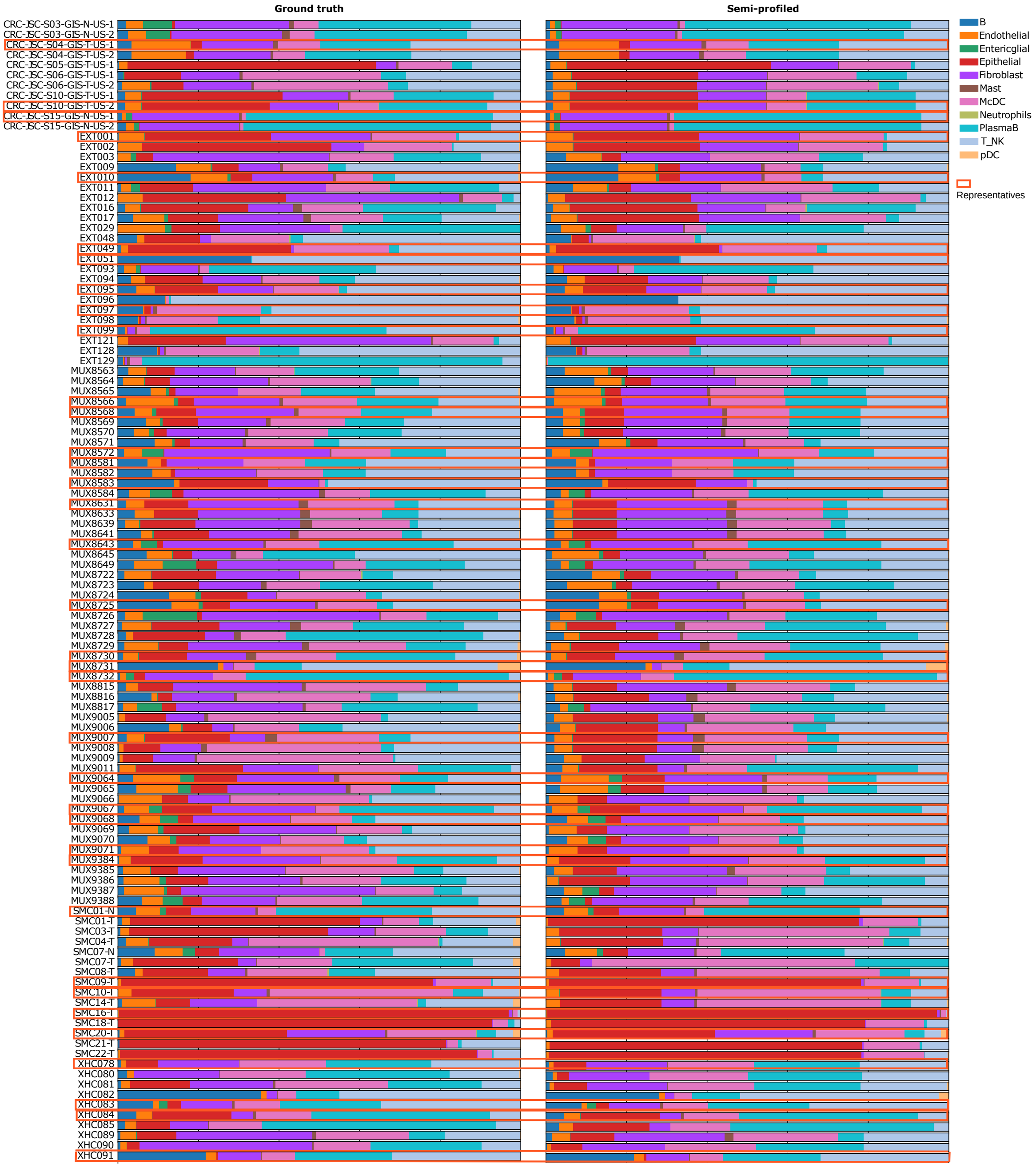


**Fig. S29**: **Deconvolution results for every sample in the colorectal cancer cohort.** The average Pearson Correlation between the real and semi-profiled versions is 0.928, and the RMSE is 0.118.

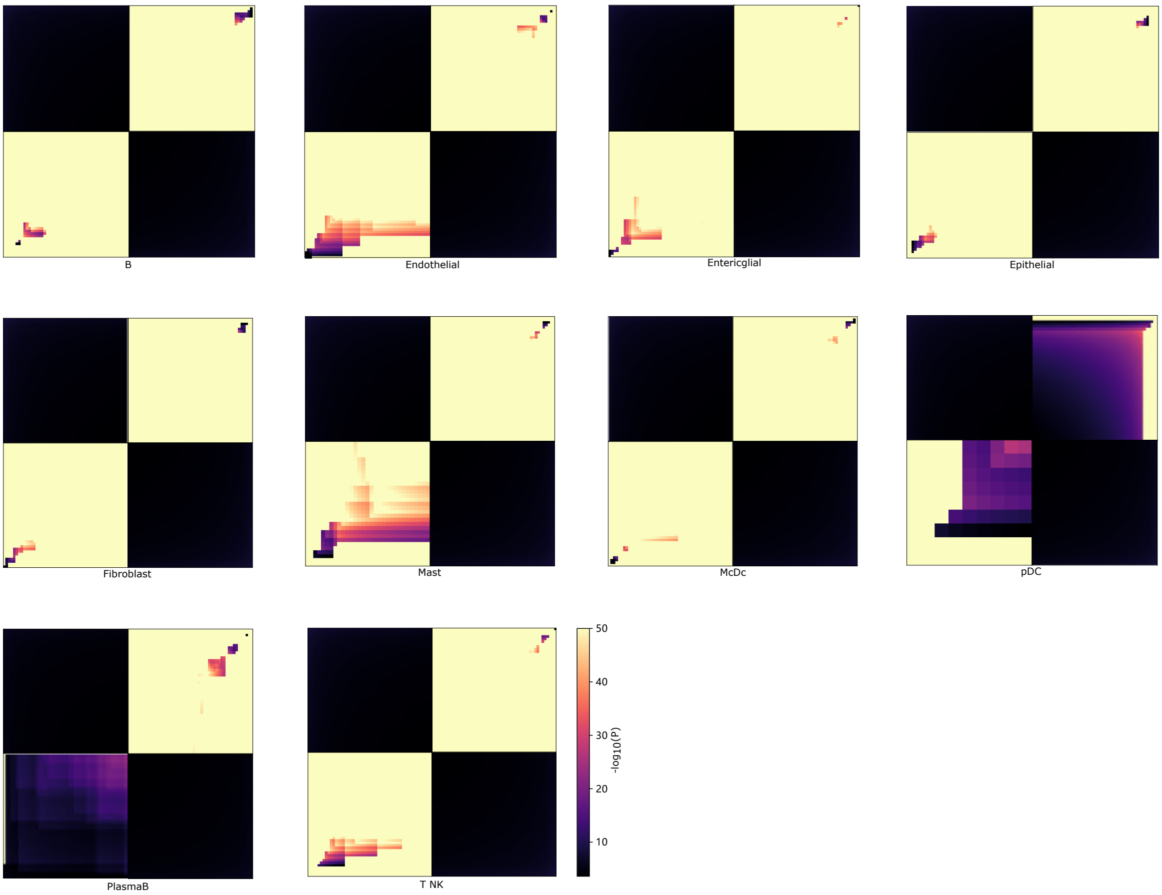


**Fig. S30**: **The RRHO plots for other cell types in the colorectal cancer dataset.** The top 50 positive and negative markers, identified using real-profiled and semi-profiled datasets, are utilized for the plots.

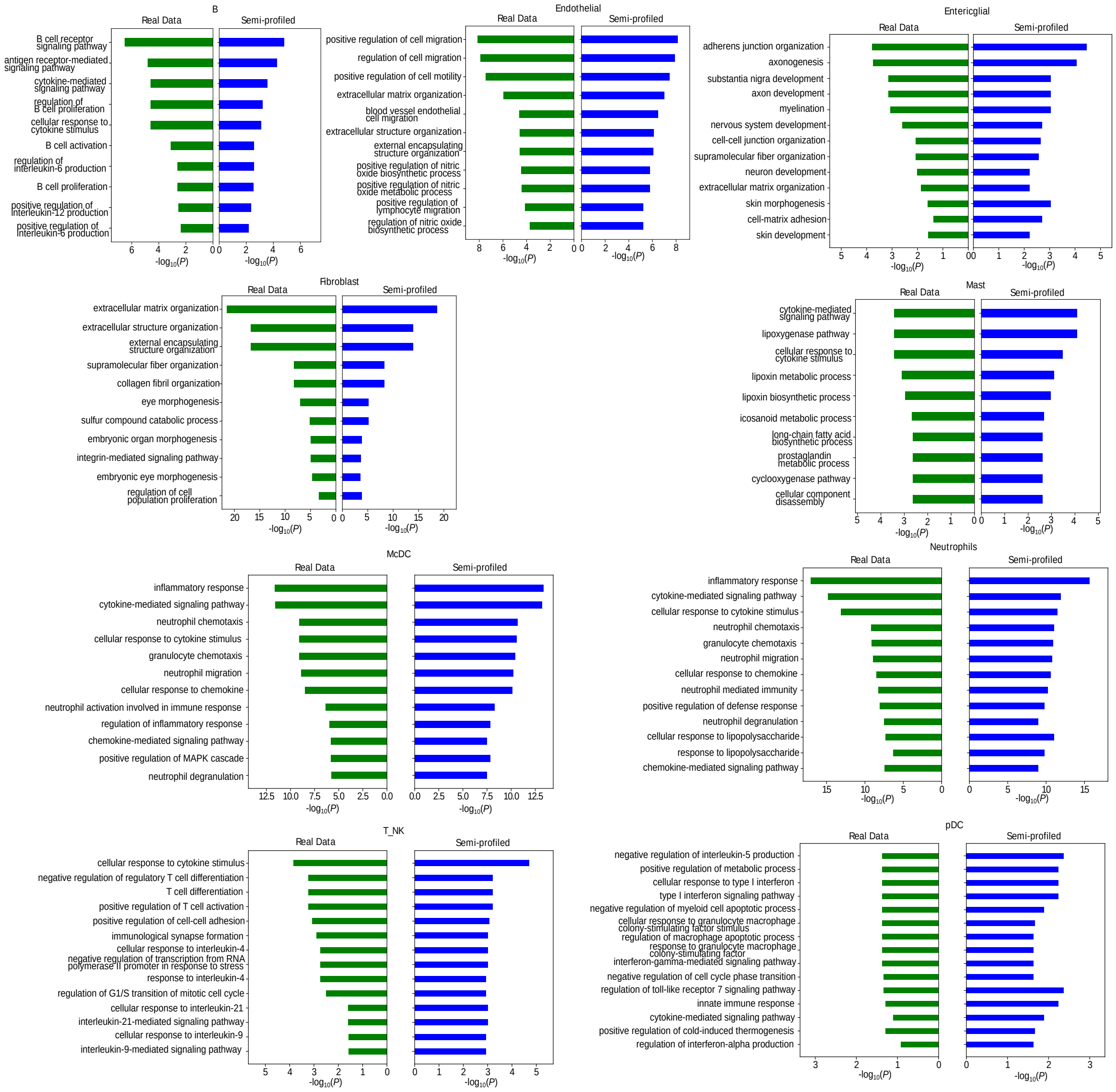


**Fig. S31**: **GO enrichment analysis for additional cell types in the colorectal cancer dataset.** The top 100 signature genes are used for the analysis. The Pearson Correlations between the negative logged p-values (bar lengths) in the real-profiled and semi-profiled versions for each cell type are as follows: B: 0.940, Endothelial: 0.969, Entericglial: 0.806, Fibroblast: 0.989, Mast: 0.946, McDC: 0.995, Neutrophils: 0.870, pDC: 0.245, T_NK: 0.603.

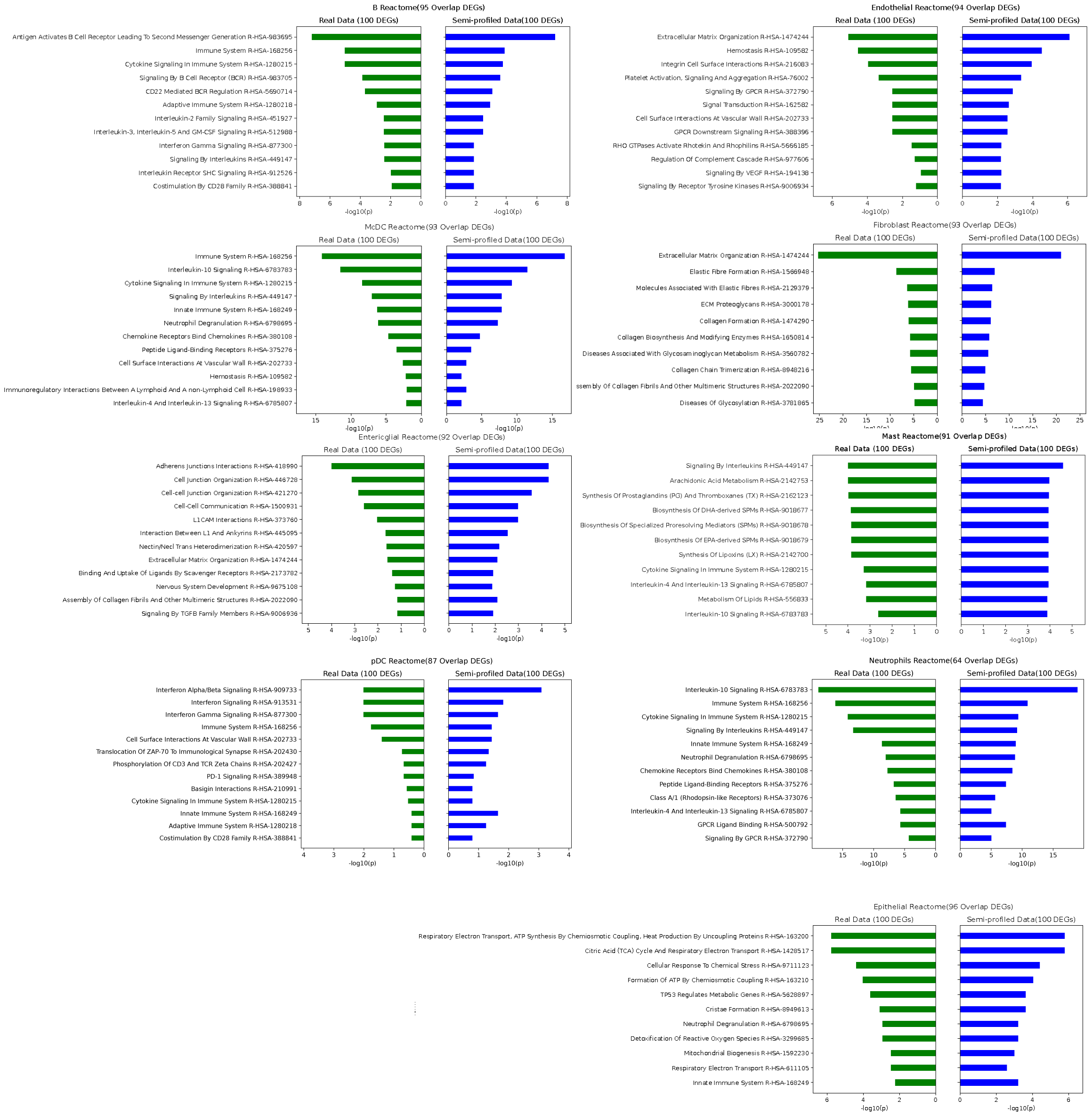


**Fig. S32**: **Reactome enrichment analysis for additional cell types in the colorectal cancer dataset.** The top 100 signature genes are used for the analysis. The Pearson Correlations between the negative logged p-values (bar lengths) in the real-profiled and semi-profiled versions for each cell type are as follows: B: 0.959, Endothelial: 0.929, Entericglial: 0.959, Epithelial: 0.996, Fibroblast: 0.997, Mast: 0.368, McDC: 0.990, Neutrophils: 0.864, pDC: 0.697, T_NK: 0.912.

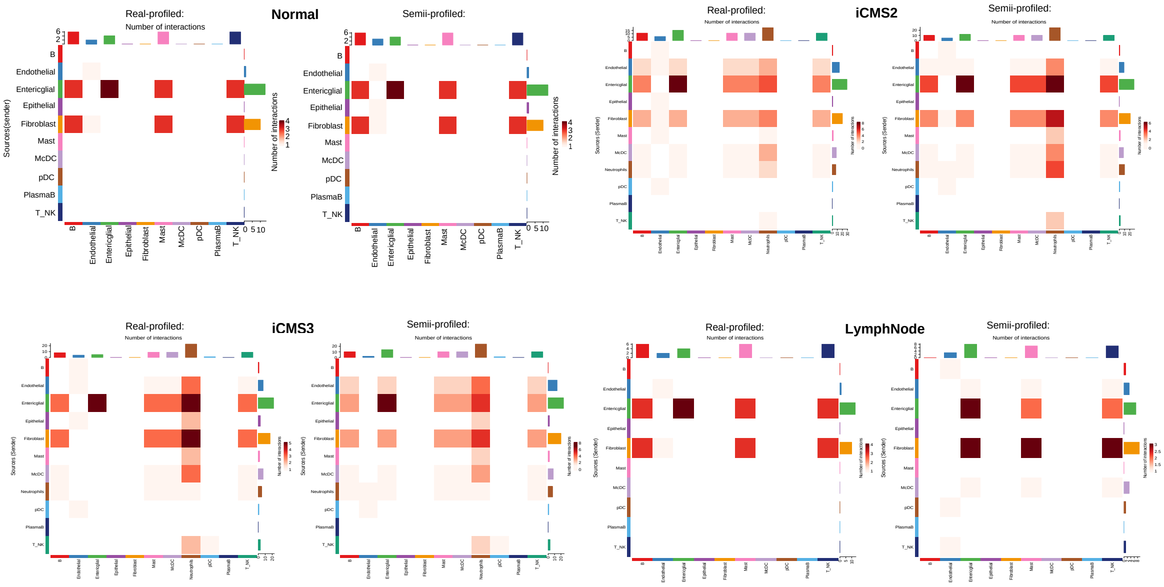


**Fig. S33**: **Cell-cell interaction analysis results comparison for all tissue types in the colorectal cancer dataset.** The Pearson Correlations between the real and semi-profiled versions of each tissue type are: Normal: 0.993, LymphNode: 0.700, iCMS2: 0.953, iCMS3: 0.922.

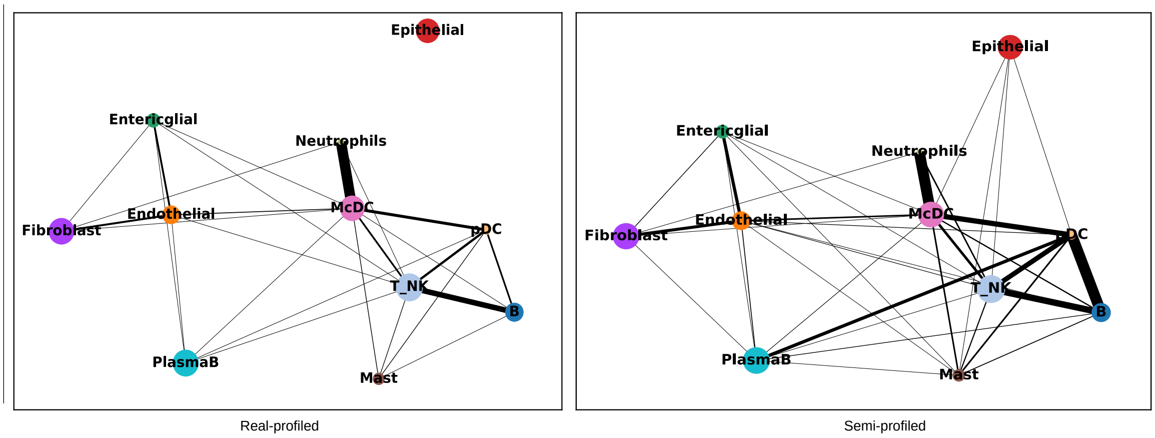


**Fig. S34**: **PAGA results comparison for the colorectal cancer dataset.** The Pearson Correlation between the two connectivity matrices is 0.830. For easier comparison, the positions of the nodes are synchronized between the ground truth and the semi-profiled PAGA results.

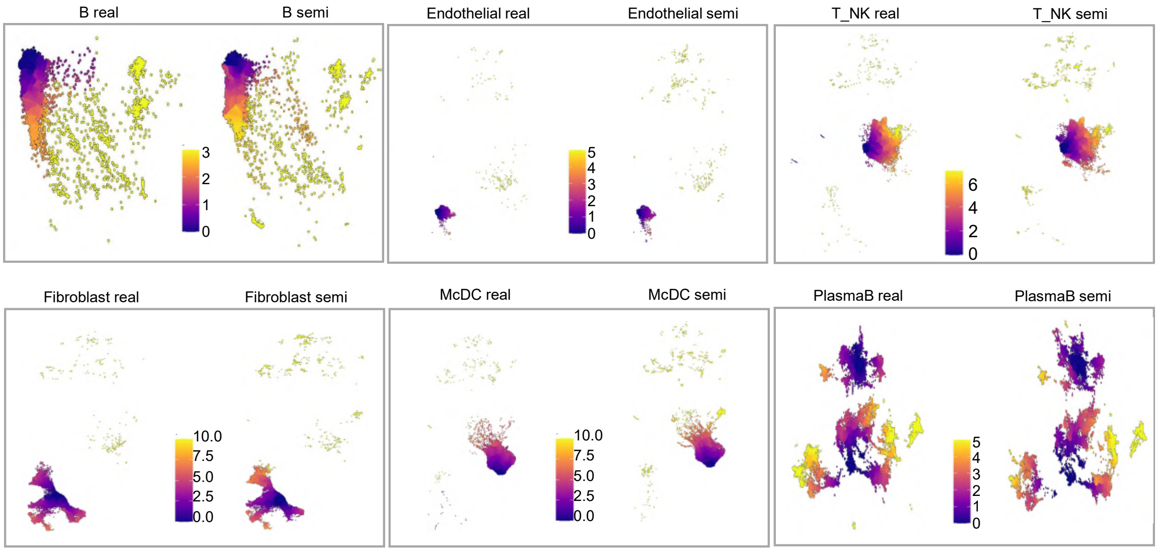


**Fig. S35**: **Pseudotime analysis results comparison for other cell types in the colorectal cancer dataset.** Cell types with insufficient cells to form a cluster structure were excluded.

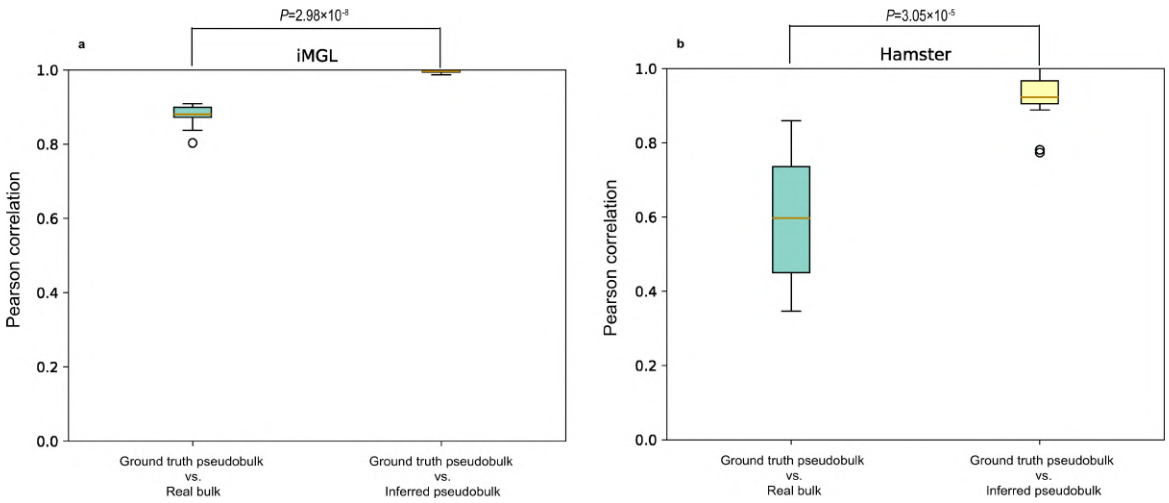


**Fig. S36**: **Comparative analysis of similarity of inferred pseudobulk and real bulk against ground truth pseudobulk**. This figure contrasts the ground truth pseudobulk data against both real bulk data and inferred pseudobulk data using our strategy across two datasets featuring paired single-cell and real bulk data. The results illustrate a significantly higher similarity of our inferred pseudobulk data (based on the real bulk) to the ground truth pseudobulk (of the target single-cell data) compared to the real bulk data (direct usage), as confirmed by p-values derived from Wilcoxon tests. **a**, iMGL cohort results comparison between inferred pseudobulk and real bulk in terms of their concordance with ground truth pseudobulk. **b**, Hamster cohort results comparison between inferred pseudobulk and real bulk in terms of their concordance with ground truth pseudobulk.

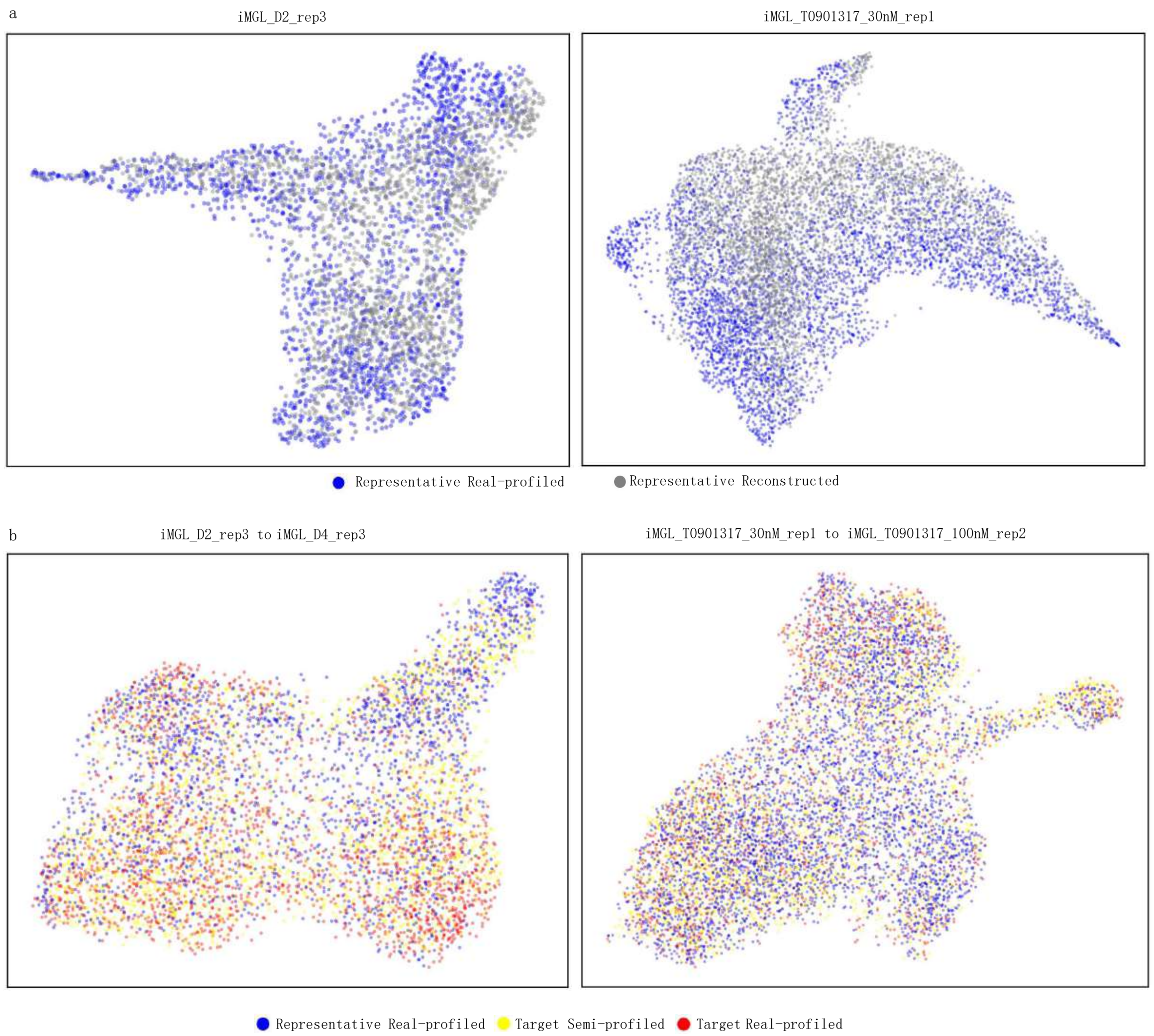


**Fig. S37**: **Examples of the in silico single-cell data inference for target samples in the iMGL cohort.** The first row shows the reconstruction of the corresponding representatives. The second row shows the cell distribution of the representative, inferred target sample, and target sample ground truth.

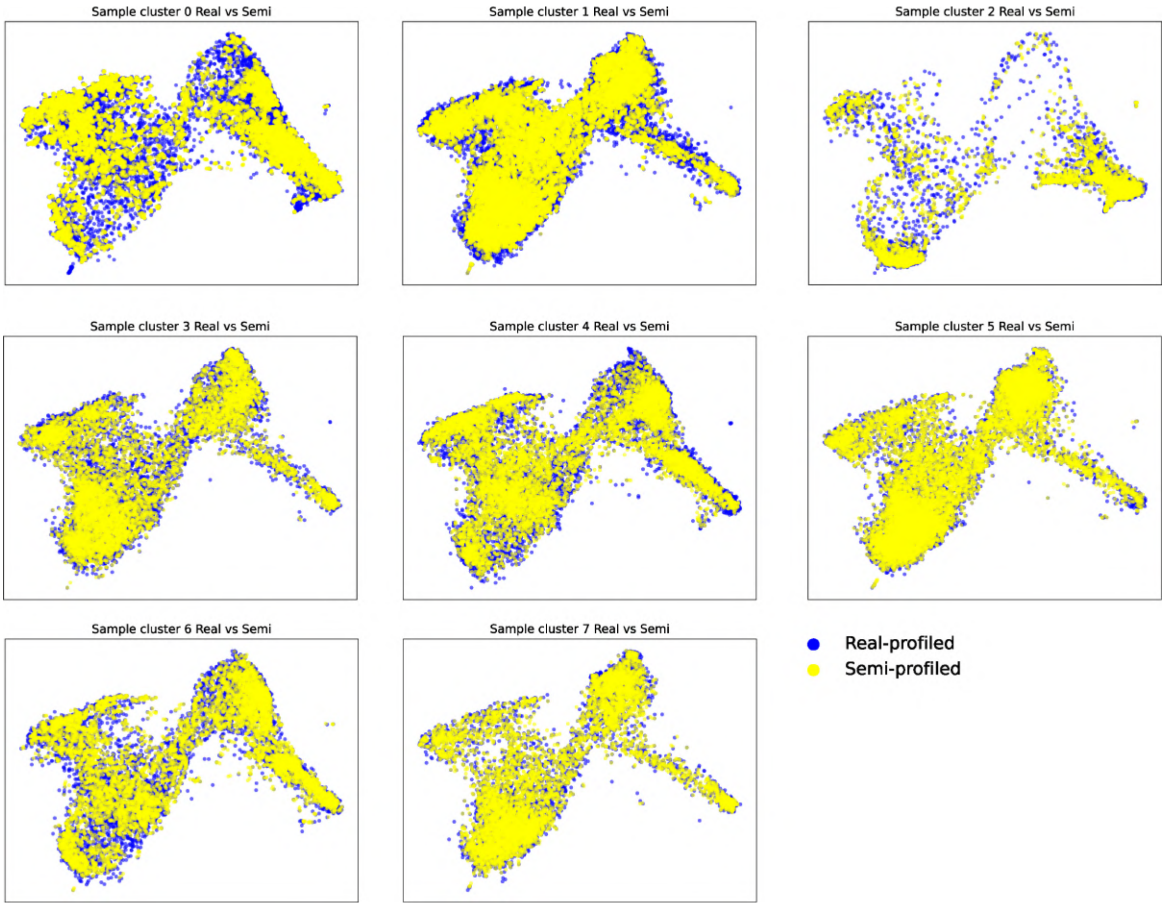


**Fig. S38**: **iMGL real-profiled and semi-profiled dataset comparison separated by sample clusters.**

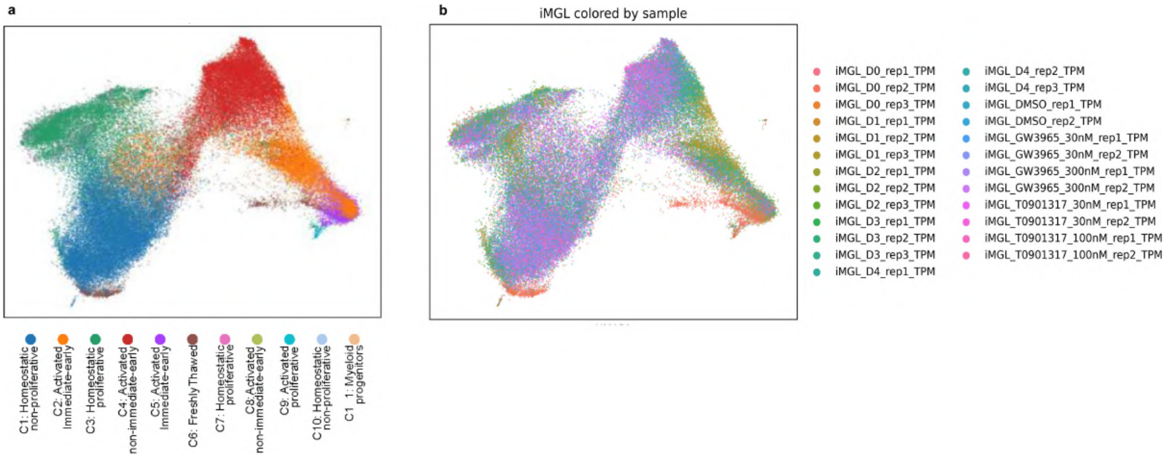


**Fig. S39**: **UMAP visualizing the cell distribution difference between samples in the iMGL dataset. a**, UMAP of the real-profiled IMGL dataset colored by cell types. **b-e**, UMAPs of the real-profiled iMGL dataset colored by samples.

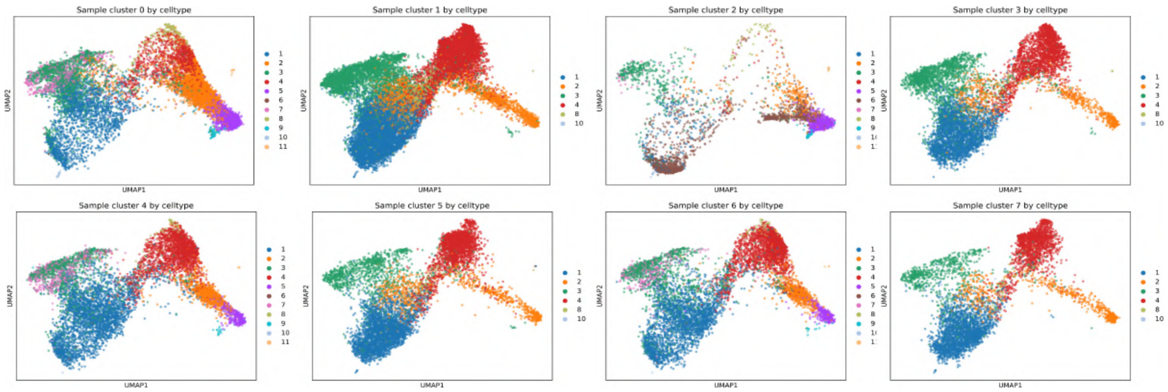


**Fig. S40**: **UMAP visualization of each iMGL sample cluster showing different cell type distributions in different sample clusters.**

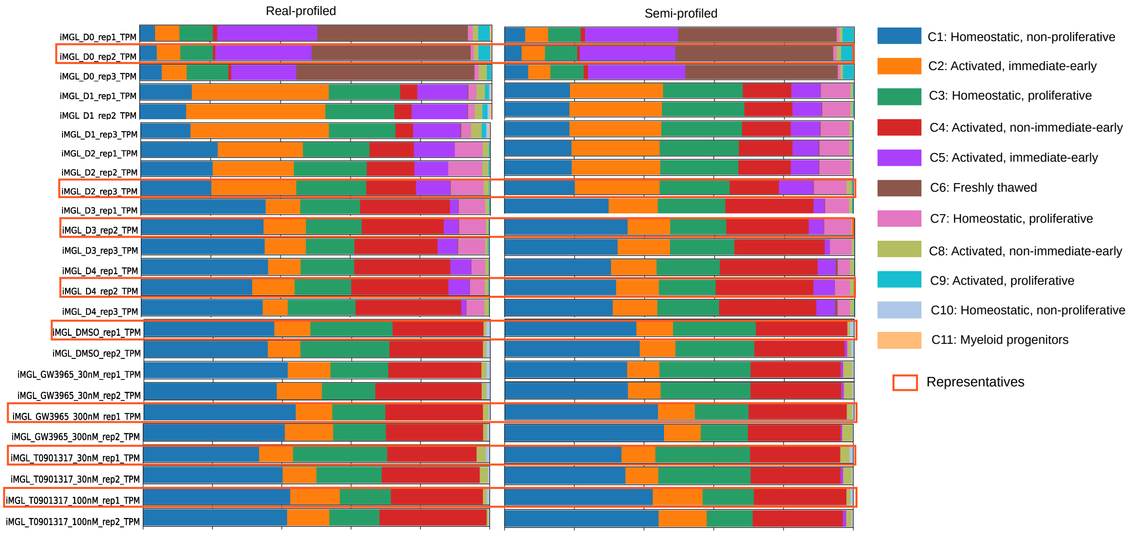


**Fig. S41**: **Deconvolution results for every sample in the iMGL cohort.** The average Pearson Correlation is 0.973, and the RMSE is 0.069.

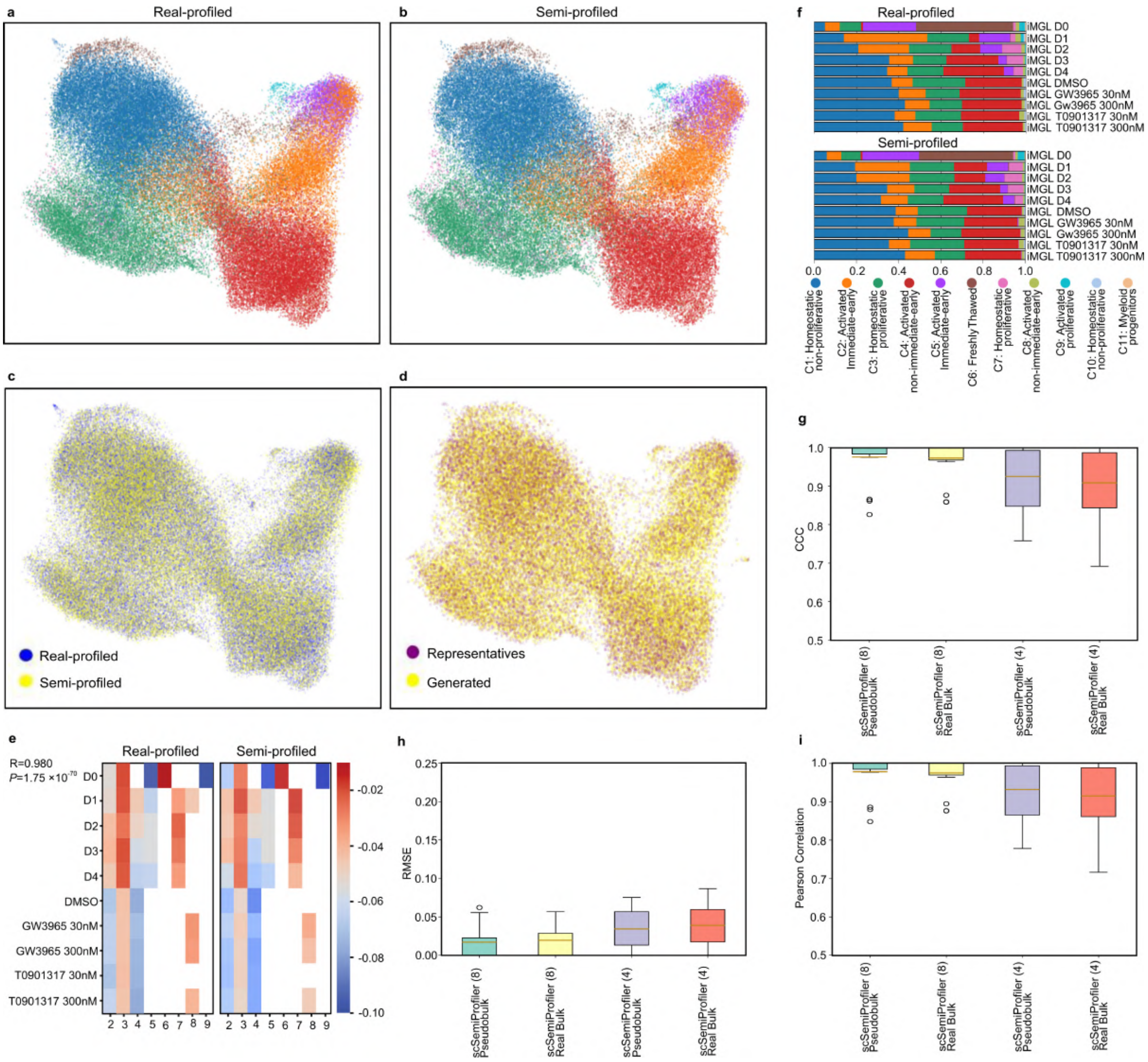

**Fig. S42**: **Comparing semi-profiled iMGL dataset generated based on pseudobulk data and real-profiled ground truth.** Please note this pseudobulk is not inferred from the real bulk. Instead, it is a direct average of the single-cell data. **a**, Real-profiled iMGL data colored by cell type. Colors are consistent with (**b**) and (**f**). **b**, Semi-profiled iMGL data based on pseudobulk colored by cell type. **c**, Combined UMAP comparing real-profiled and pseudobulk-based semi-profiled IMGL dataset. **d**, Pseudobulk-based semi-profiled IMGL dataset colored by cell origins (from representative samples or generated by the deep generative model). **e**, "activation of immune response" GO term activation pattern comparison. **f**, Comparing cell type proportions under different experimental conditions between datasets. The Pearson correlation between the real-profiled and pseudobulk-based semi-profiled versions of cell type proportions under different conditions are: iMGL_D0: 0.998, iMGL_D1: 0.872, iMGL_D2: 0.997, iMGL_D3: 0.998, iMGL_D4: 0.995, iMGL_DMSO: 0.998, iMGL_GW_30: 0.983, iMGL_GW_300: 0.999, iMGL_T_30: 0.993, iMGL_T_300: 0.999. **g,h,i**, Deconvolution performance comparison between pseudobulk-based and real bulk-based semi-profiling using CCC, RMSE, Pearson correlation respectively.

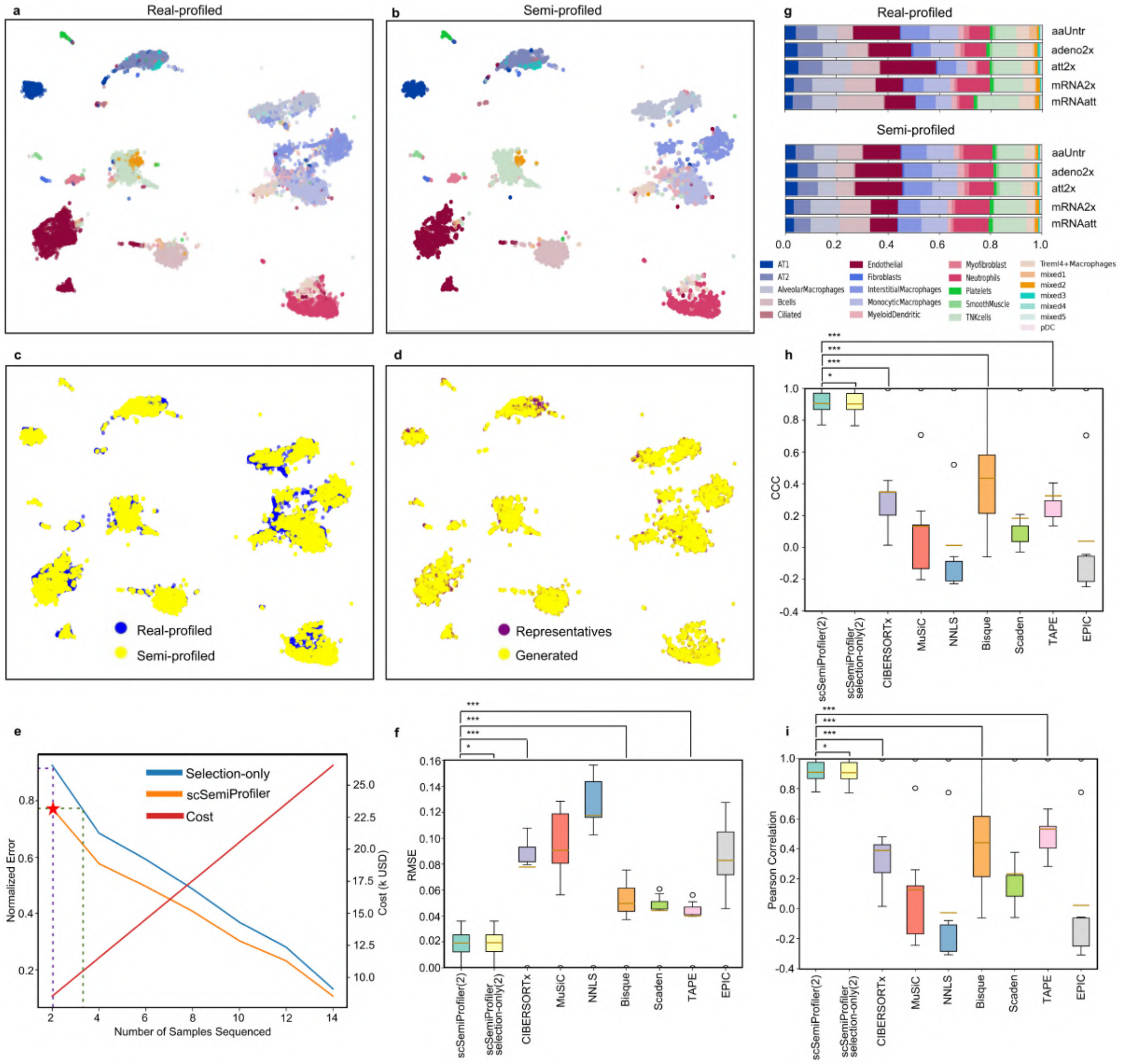


**Fig. S43**: **Comparing real-profiled and semi-profiled hamster dataset.** **a**, UMAP visualization of cells in the real-profiled hamster cohort, with colors representing cell types. Colors are consistent with (**g**). **b**, UMAP visualization of the semi-profiled hamster dataset. **c**, Jointly visualize real-profiled and semi-profiled hamster cohorts, showing a high similarity between them. **d**, UMAP of the semi-profiled cohort, with colors representing cells from the 2 representative samples or from the rest 14 inferred non-representative samples. **e**, Error trajectory of *scSemiProfiler* as more representatives are selected. **f, h, i**, Deconvolution performance benchmarking using RMSE, CCC, and Pearson correlation respectively. All results show a significant lead of our method over other deconvolution methods. **g**, Stacked bar plots visualizing the cell type proportions in different experiment treatments in the real-profiled and semi-profiled cohorts. The Pearson correlations between the real-profiled and semi-profiled cohorts under different treatments are: aaUntr: 0.934, adeno2x: 0.963, att2x: 0.898, mRNA2x: 0.993, mRNAatt: 0.873.

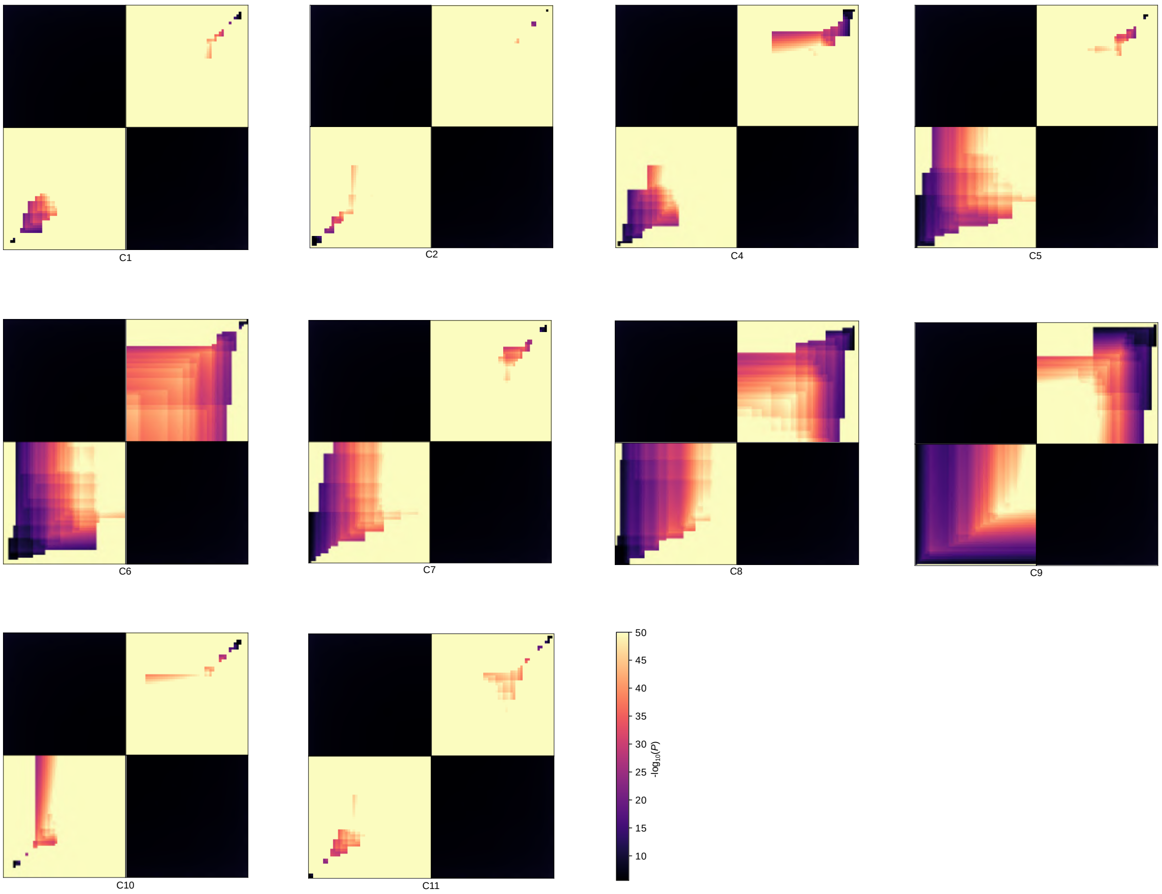


**Fig. S44**: **The RRHO plots for other cell types in the iMGL dataset.** The top 50 positive and negative markers, identified using real-profiled and semi-profiled datasets, are utilized for the plots.

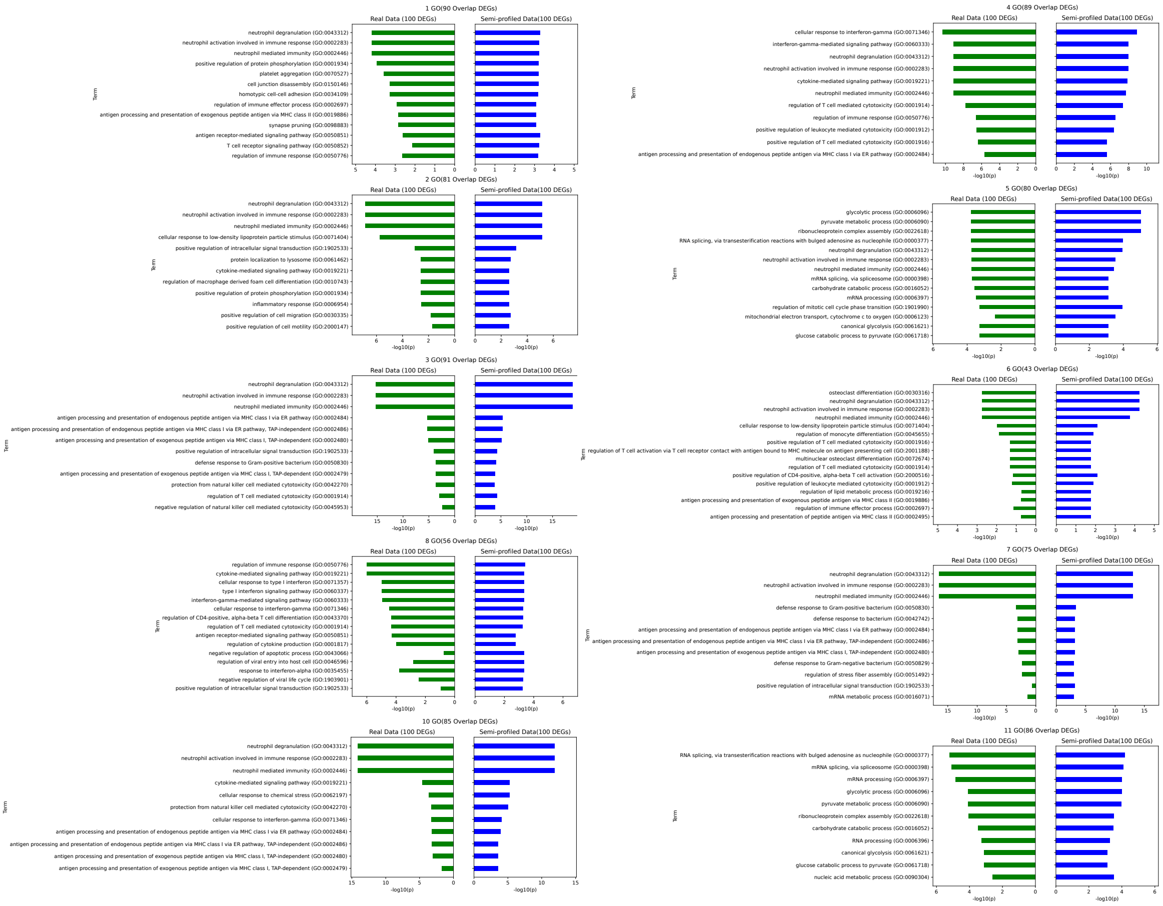


**Fig. S45**: **GO enrichment analysis for additional cell types in the iMGL dataset.** The top 100 signature genes are used for the analysis. Cell types without at least 10 GO terms are not included. The Pearson Correlations between the significance level in real-profiled and semi-profiled version for each cell type are: C1 Homeostatic, non-proliferative: 0.352, C2 Activated, immediate-early: 0.978, C3 Homeostatic, proliferative: 0.995, C4 Activated, non-immediate-early: 0.974, C5 Activated, immediate-early: 0.353, C6 Freshly thawed: 0.909, C7 Homeostatic, proliferative: 0.993, C8 Activated, non-immediate-early: 0.04, C9 Activated, proliferative: 0.992, C11 Myloidprogenitors: 0.855

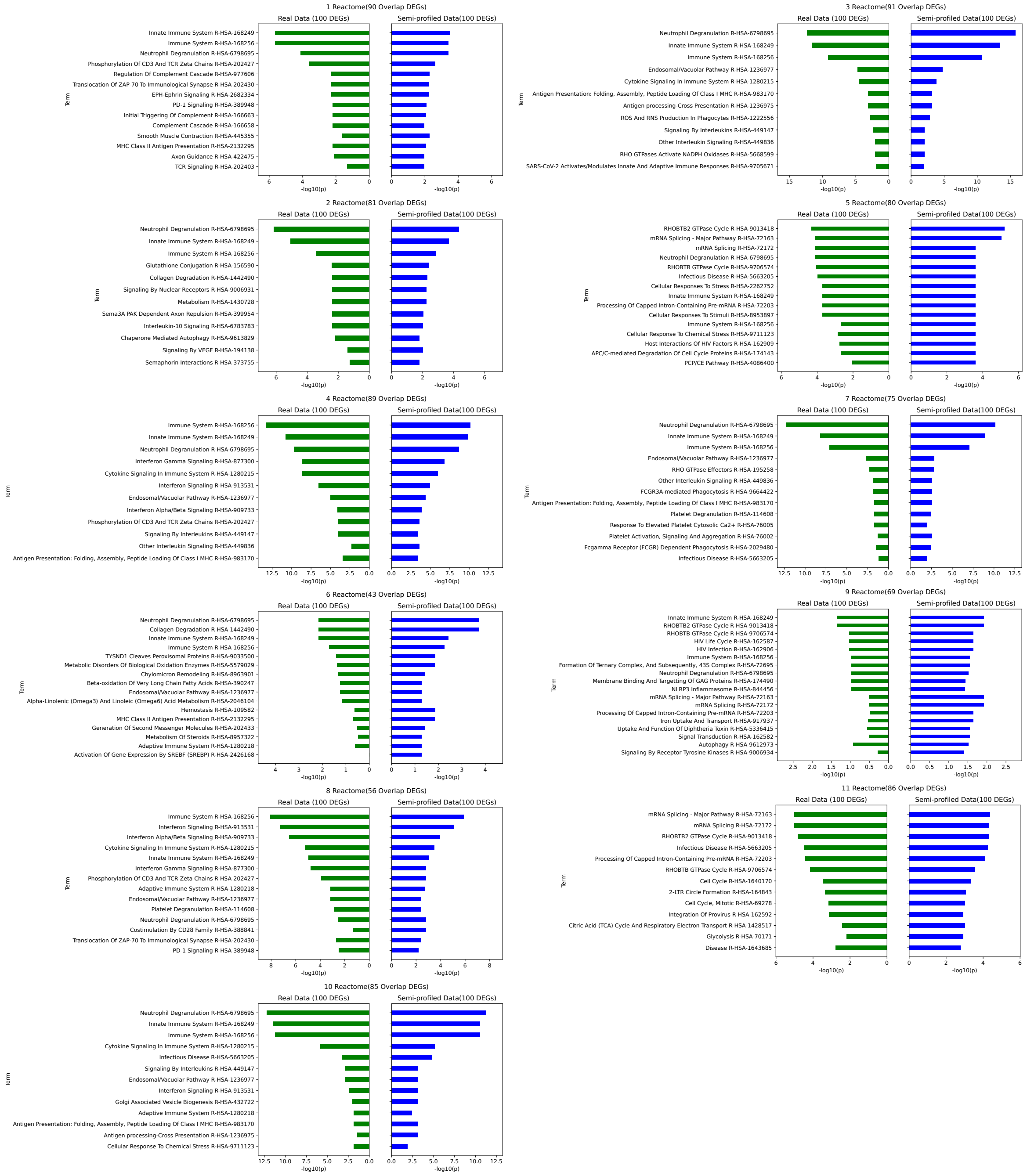


**Fig. S46**: **Reactome enrichment analysis for other cell types in the iMGL dataset.** The top 100 signature genes are used for the analysis. The Pearson Correlations between the significance level in real-profiled and semi-profiled version for each cell type are: C1 Homeostatic, non-proliferative: 0.940, C2 Activated, immediate-early: 0.974, C3 Homeostatic, proliferative: 0.995, C4 Activated, non-immediate-early: 0.961, C5 Activated, immediate-early: 0.414, C6 Freshly thawed: 0.755, C7 Homeostatic, proliferative: 0.984, C8 Activated, non-immediate-early: 0.889, C9 Activated, proliferative: 0.199, C10 Homeostatic, non-proliferative: 0.987, C11 Myeloidprogenitors: 0.943

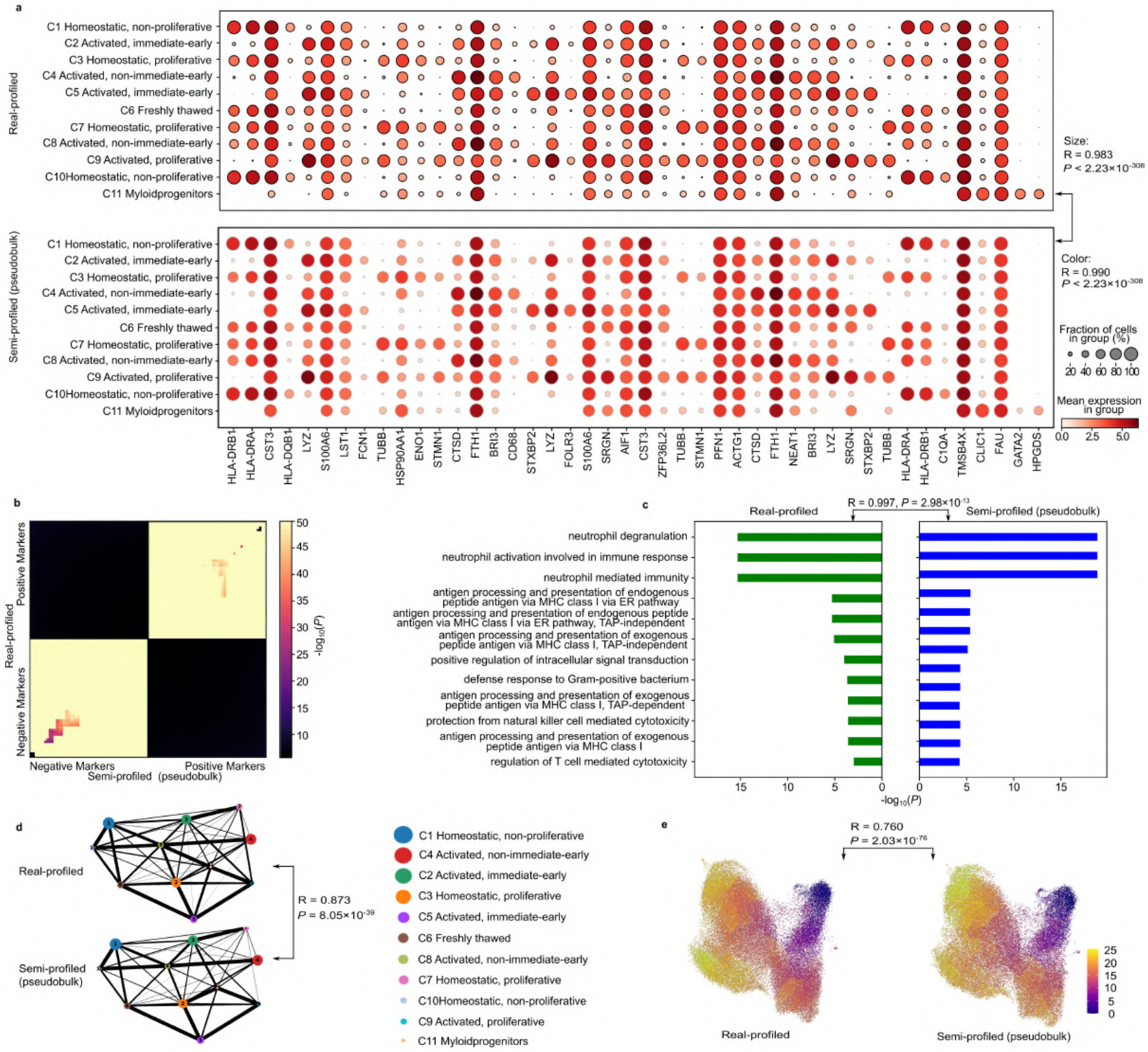

**Fig. S47**: **iMGL real-profiled cohort and pseudobulk-based semi-profiled cohort downstream single-cell analysis results comparison.** Please note this pseudobulk is not inferred from the real bulk. Instead, it is a direct average of the single-cell data. **a**, Dot plots visualizing the cell type signature genes. **b**, RRHO plots comparing the cell type biomarkers in two datasets. **c**, C3 Cell type markers GO enrichment analysis results comparison. **d**, PAGA results generated using two versions of datasets comparison. **e**, Pseudotime analysis results comparison.

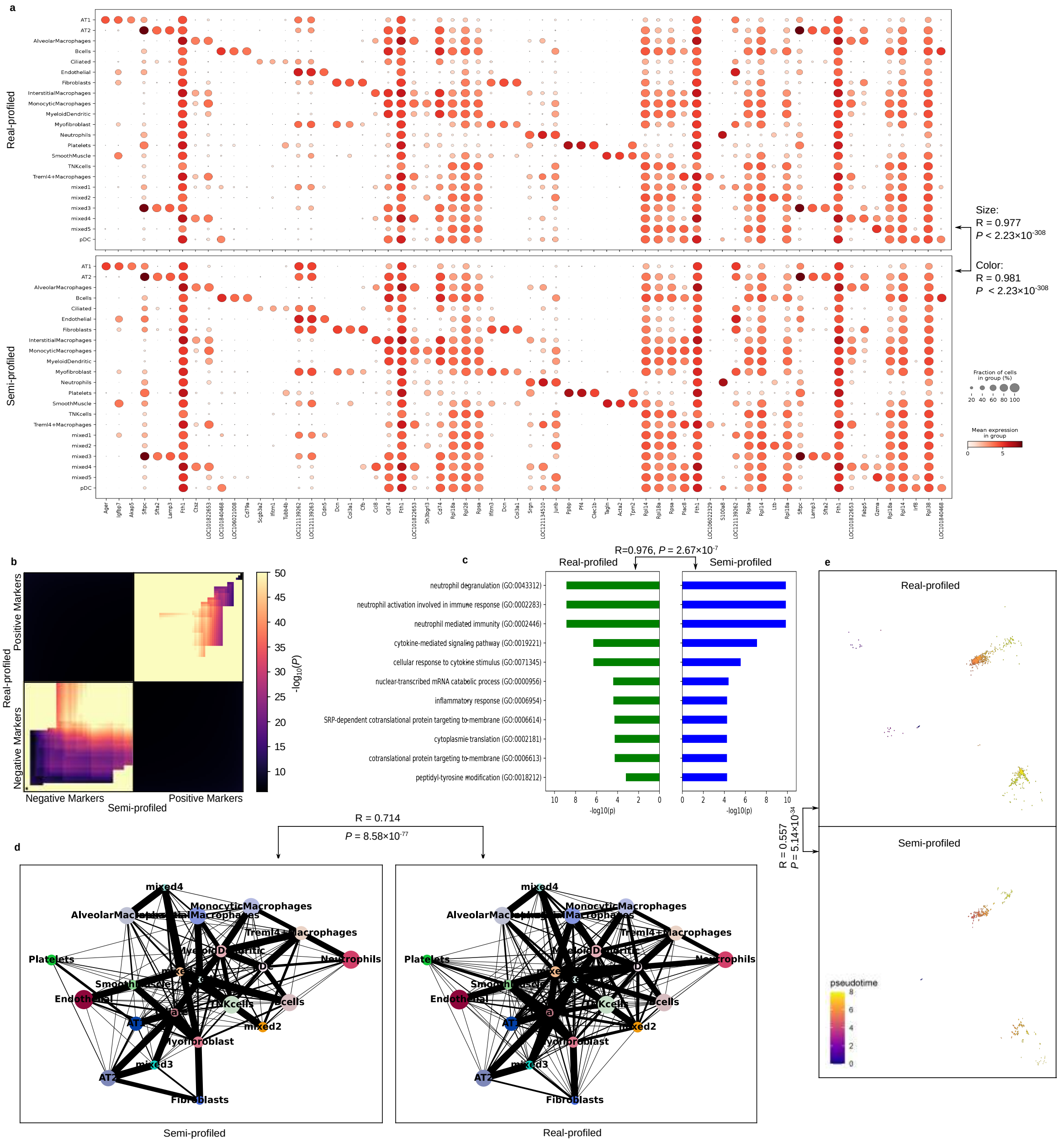


**Fig. S48**: **Comparing the downstream analysis results using the semi-profiled and real-profiled hamster cohorts.** **a**, Dot plots showing similar cell type signature genes expression patterns in the semi-profiled and real-profiled cohorts. **b**, RRHO plot comparing the Treml4+Macrophages markers found using the two versions of datasets. **c**, Treml4+Macrophages cell type signature genes GO enrichment analysis results comparison. **d**, PAGA analysis results generated from the two versions of hamster datasets show high similarity. **e**, Pseudotime results comparison using the real-profiled and *in silico* generated Treml4+Macrophages.

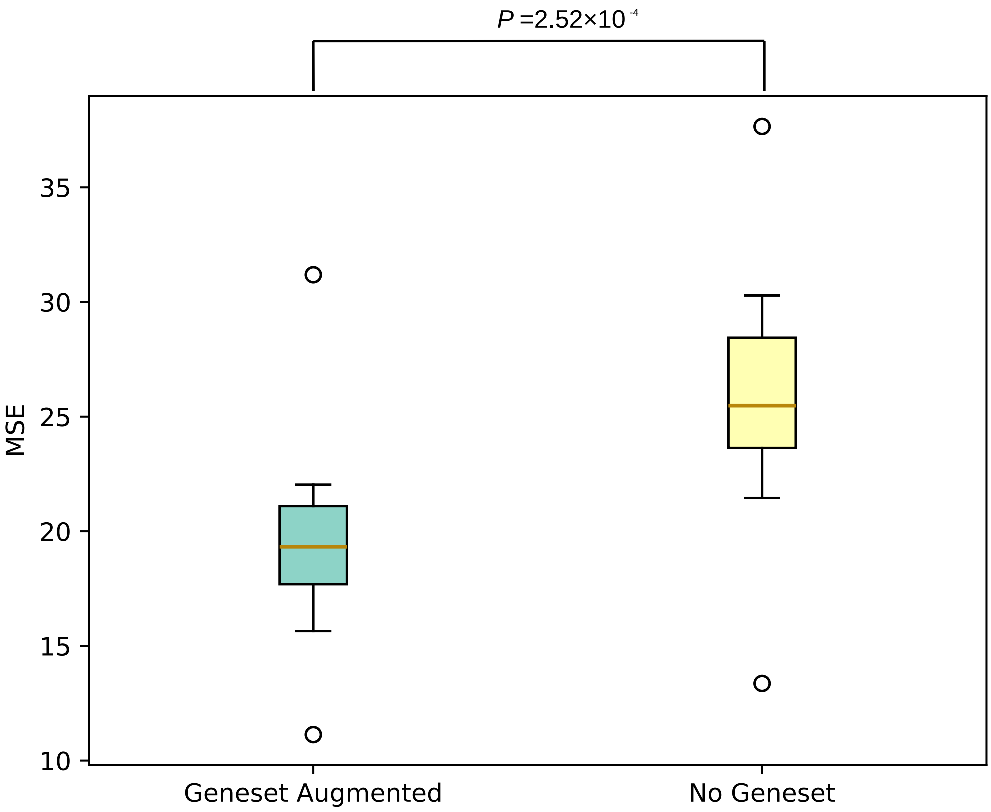


**Fig. S49**: **Gene set score ablation study.** With gene set scores augmented, our generator can perform more accurate reconstructions on the cortex dataset (PMID: 25700174) with lower average MSE (19.33 vs. 25.48, Wilcoxon test p-value: $2.52 \times 10^{-4}$).

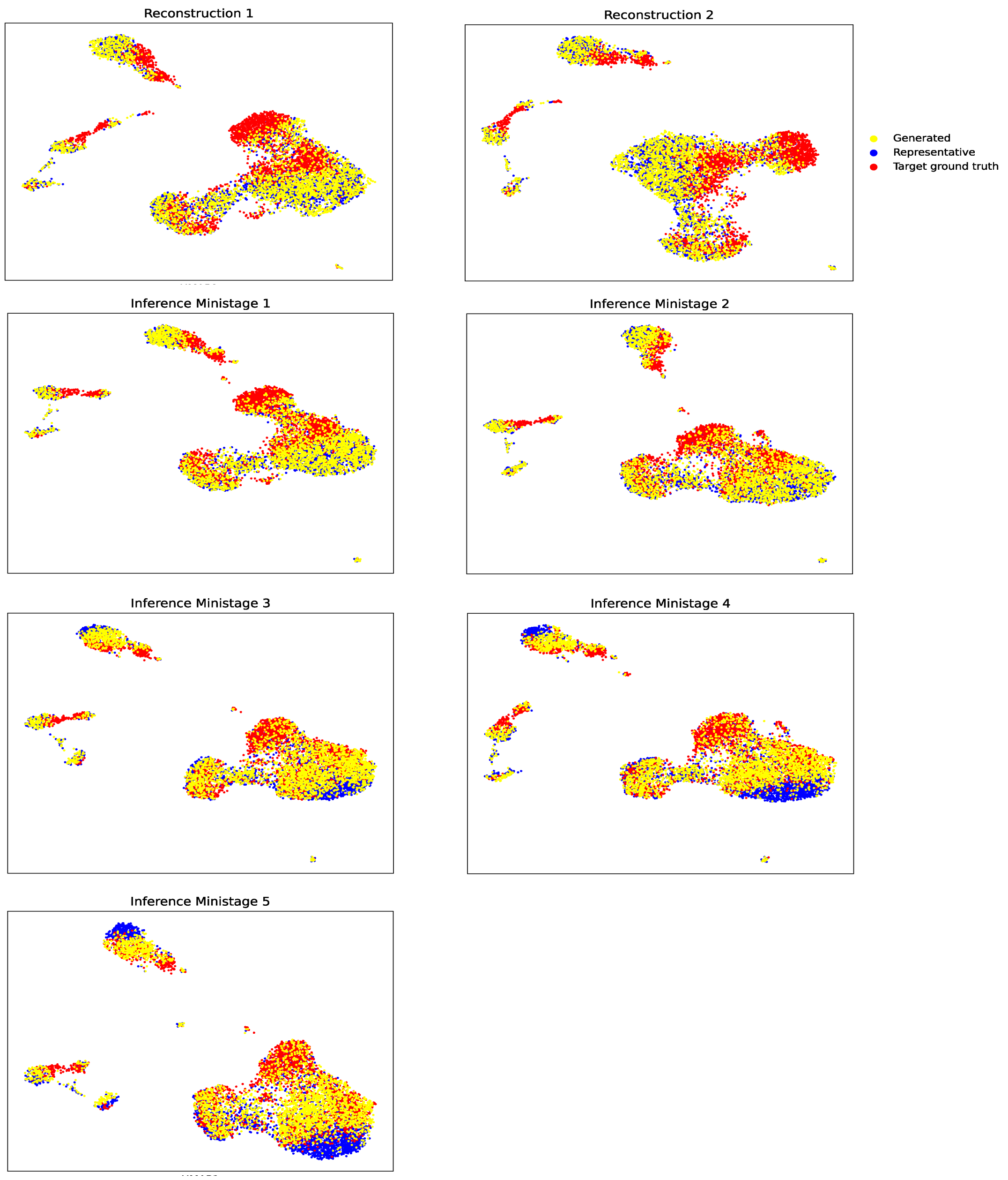


**Fig. S50**: **Visualization of the target *in silico* inference training process in mini-stages**. UMAP visualizations of real-profiled target sample's cells (red), real-profiled representative cells (blue), and *in silico generated cells (yellow)*. With successive training stages, the *in silico* generated cells increasingly resemble the target ground truth cells.

**Algorithm 1** Semi-profiling using scSemiProfiler

```
Input: Budget
Output: Semi-profiled single-cell data for the entire cohort
Main Routine:
Perform bulk sequencing of all samples.
Selected representatives ← InitialRepresentativeSelection(Bulk Data)
Conduct single-cell sequencing on selected representatives.
Pretrained models ← RepresentativeReconstruction(Selected single-cell data, Bulk Data)
Inferred single-cell data ← InSilicoInference(All bulk data, Selected single-cell data, Pretrained models)
Next representatives ← ActiveLearning(Bulk data, Current semi-profiled single-cell cohort)
If stopping criteria are not met, repeat from "Conduct single-cell sequencing on selected representatives."
return Semi-profiled single-cell data for the entire cohort.

procedure InitialRepresentativeSelection(Bulk Data)
    Cluster samples based on bulk data.
    Select a representative sample from each cluster.
    return List of selected representatives.
end procedure

procedure RepresentativeReconstruction(Selected single-cell data, Bulk Data)
    Initialize Pretrained models
    For each selected representative:
        Train a VAE generator to reconstruct single-cell data (Pretrain 1.1).
        Jointly train generator and discriminator (Pretrain 1.2).
        Enhance generator training in full batch mode with added bulk data loss (Pretrain 2.1).
        Similar to Pretrain 2.1, but train discriminator jointly (Pretrain 2.2).
    return Pretrained models for selected representatives.
end procedure

procedure InSilicoInference(All bulk data, Selected single-cell data, Pretrained models)
    Initialize inferred single-cell data
    For each non-representative sample:
        Select the pretrained model of the closest selected representative.
        Fine-tune the model incorporating bulk sample differences.
        Conduct mini-stages 1-5 for tuning gene expression accuracy.
        Infer single-cell profile for the non-representative sample.
    return Inferred single-cell data for all non-representative samples.
end procedure

procedure ActiveLearning(Bulk data, Current semi-profiled single-cell cohort)
    Analyze heterogeneity within clusters using current semi-profiled single-cell cohort and the bulk data
    Select next round of representatives to minimize heterogeneity.
    Update cluster membership.
    return List of next representatives.
end procedure
```

1

**Fig. S51**: **Pseudocode of *scSemiProfiler*'s pipeline.**
